# Guard cell size and initial conductance influence stomatal closure kinetics

**DOI:** 10.64898/2026.05.17.725794

**Authors:** Christopher D. Muir, Wei Shen Lim

## Abstract

- In fluctuating environments, the kinetics of stomatal opening and closing influence the balance between carbon gain and water loss. Smaller guard cells may respond faster to fluctuating environmental conditions because of their greater surface area for osmolyte flux relative to cell volume. A related hypothesis is that operational stomatal conductance (*g*_op_) is often well below its theoretical maximum (*g*_max_) because at this stomatal aperture, guard cell volume is poised to change rapidly with small changes in turgor pressure.
- We analyzed 2,125 estimates of stomatal closure kinetics in response to an abrupt increase in vapor pressure deficit (VPD) among 29 diverse wild tomato populations in the genus *Solanum*.
- Leaves with small guard cells and a lower initial stomatal conductance (*g*_i_) closed faster, but each explained variation in kinetic parameters at different levels of biological organization. Guard cell size had high phylogenetic heritability and varied relatively little within populations, whereas *g*_i_ varied mostly among individuals and between light intensity treatments.
- Smaller stomata can be speedier, but the current physiological state of the leaf also affects the responsiveness of stomatal conductance to fluctuations. Selection on stomatal speed may influence not only anatomical traits like guard cell size, but also physiological controls on *g*_op_.

## Introduction

In nature, change is the only constant. One way that vascular plants (tracheophytes) cope with change is by adjusting stomatal pore aperture to optimize the trade-off between carbon gain and water loss (Cowan & Farquhar, 1977; Raven, 2002; Hetherington & Woodward, 2003). In the absence of physical limits and energetic tradeoffs, natural selection would favor plants that could instantaneously adjust stomatal conductance (*g*_sw_) to perfectly track dynamic environmental conditions (Lawson *et al*., 2010). In reality, stomatal responses take time, creating a lag between actual and optimal stomatal aperture. For example, many species use hydroactive feedback mechanisms to precisely control stomatal responses to changing vapor pressure deficit (VPD), the driving force for evaporation (Buckley, 2019), but hydroactive control takes time. Leaves must sense a change in VPD through as-yet-unknown mechanisms and then transduce a signal that causes efflux of osmotic solutes to alter guard cell turgor more than the surrounding epidermal cells. Abscisic acid (ABA) dependent responses require several minutes to accumulate or synthesize hormone pools *de novo* (Xie *et al*., 2006; Bauer *et al*., 2013; McAdam *et al*., 2016; Desai & Stroock, 2025; Pichaco *et al*., 2026); ABA-independent stomatal responses require changes in gene expression and protein synthesis that take time (Jalakas *et al*., 2021). Once anion channels are activated, the rate of change in stomatal aperture depends on the rate of change in guard cell turgor (*dP*_g_/*dt*) and the sensitivity of aperture to change in turgor (*da*_S_/*dP*_g_). As we discuss below, these two factors (*dP*_g_/*dt* and *da*_S_/*dP*_g_) may be influenced by guard cell size and the ratio of operational to maximum stomatal conductance (*f*_gmax_), respectively.

Over a century ago Darwin (1898) observed that after an initial increase, leaf transpiration decreases in response to decreasing humidity. He correctly attributed changes in transpiration to changes in stomatal aperture and subsequent studies have confirmed that stomatal aperture responds to humidity (Lange *et al*., 1971) and, more specifically, transpiration rate (Mott & Parkhurst, 1991; Monteith, 1995), which is the product of VPD and *g*_sw_. The VPD is related to relative humidity (RH) but also temperature. In angiosperms and some pteridophytes, a sudden increase in VPD, usually induced by lower RH at a constant temperature, causes a transient increase in *g*_sw_, as Darwin observed (Cowan, 1972; Buckley *et al*., 2011; Westbrook & McAdam, 2021). After a period of several minutes, stomata close until the leaf reaches a new steady-state *g*_sw_. These phases of stomatal response to VPD are often referred to as the “wrong-way response” (WWR) and “right-way response” (RWR), respectively. The normative ‘wrong’ and ‘right’ appellations refer to the presumed adaptive significance. At steady state, *g*_sw_ is negatively correlated with VPD, all else being equal, which is interpreted as an adaptive response to the marginal carbon cost of water (Givnish & Vermeij, 1976; Cowan & Farquhar, 1977; Buckley *et al*., 2017b). Models built around this assumption generally predict steady state *g*_sw_ well, although the functional form and presumed fitness costs are areas of active research (Damour *et al*., 2010; Medlyn *et al*., 2011; Prentice *et al*., 2014; Sperry *et al*., 2017; Potkay *et al*., 2025).

Stomatal responses are as much about the journey (kinetics) as they are about the destination (steady state). If stomatal response kinetics are too slow relative to the temporal grain size of environmental variability, then it would be futile for *g*_sw_ to track environmental variation since the leaf will constantly lag the environment, perpetually fighting the last war. This may explain why *g*_sw_ can be insensitive to short sun/shade flecks (Knapp & Smith, 1990) and shaded leaves are likely primed to cope with the hydraulic demands of sudden increases in light, temperature, and VPD (Schymanski *et al*., 2013). If stomatal responses can track environmental variation, then minimizing the lag time can greatly improve resource-use efficiency (Lawson *et al*., 2010). For example, optogenetic control of stomatal aperture in response to fluctuating light increases water-use efficiency in *Arabidopsis* compared to wild-type responses (Papanatsiou *et al*., 2019). Therefore the rate of stomatal opening and closing is predicted to have a major effect on how natural selection optimizes stomatal control in variable environments (Buckley *et al*., 2023).

Much of the research on stomatal kinetics focuses on light responses because light intensity is usually the most important, rapidly fluctuating environmental variable affecting photosynthesis. In nature, fluctuating light intensity simultaneously alters leaf temperature and VPD (e.g. Ishida *et al*., 1992). We focus on stomatal closure kinetics in response to an abrupt increase in VPD, rather than light, for four reasons. First of all, the primary aims and *a priori* hypotheses of this study were not to study stomatal kinetic parameters and are described in Muir *et al*. (2025). The idea to opportunistically use *g*_sw_-response curves to study stomatal kinetics came after the data had been collected, but not yet analyzed for this purpose. Second, VPD responses exhibit a similar functional form as light-responses, suggesting similar underlying mechanisms (Buckley, 2019). Factors that affect stomatal response kinetics to VPD likely generalize to other environmental variables. Third, humidity responses tend to dominate over and are faster than other environmental responses (Aasamaa & Sõber, 2011a,b), although there is variation among species (Merilo *et al*., 2014). Thus, factors that constrain maximum rates of stomatal closure are most likely to be observed in response to VPD. Finally, the ecological salience of VPD responses is increasing as climate change warms the atmosphere (Grossiord *et al*., 2020).

Fully describing stomatal responses to the environment is complex (Buckley *et al*., 2003; Jezek *et al*., 2021; Desai & Stroock, 2025), but the dynamics of *g*_sw_ in response to an abrupt environmental change is well described by relatively simple phenomenological functions (Vico *et al*., 2011; McAusland *et al*., 2016). A four-parameter equation describes stomatal response kinetics to a step change in light intensity across dozens of plant species (Woning *et al*., 2026):

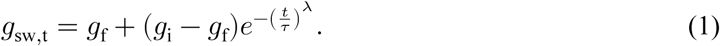

In this equation, *g*_sw,t_ is stomatal conductance at time *t, g*_i_ is the initial *g*_sw_ before the step change, *g*_f_ is the final *g*_sw_ after the step change, *τ* is a time constant that describes how quickly stomata respond to the step change, and *λ* is a shape parameter that describes how the response rate changes over time. When *λ*= 1, the response rate is constant over time and *τ* represents the time required for *g*_sw_ to reach 63.2% (1 − *e*^−1^) of the way between initial and final values. When *λ* > 1, the response rate increases over time, and when *λ* < 1, the response rate decreases over time. Woning *et al*. (2026) refer to *λ* as a lag-time parameter because empirically *λ* > 1. We refer to stomatal kinetic parameters *τ* (time constant) and *λ* (lag time) henceforth. The first goal of this study is to evaluate whether kinetics of the RWR after a step change in VPD are adequately described by Equation 1. If so, it suggests that some fundamentally similar mechanisms are shared between light and hydroactive stomatal responses (Grantz & Zeiger, 1986).

The main goal of this study is to test hypotheses about which stomatal anatomical traits affect kinetic parameters *τ* and *λ*. We consider two interconnected hypotheses that have thus far been treated in isolation. The first hypothesis is that smaller guard cells open and close faster because of their intrinsically greater surface area to volume ratio (Franks & Farquhar, 2007; Drake *et al*., 2013; Lawson & Vialet-Chabrand, 2019; Woning *et al*., 2026). For approximately cylindrical cell geometries, like that found in guard cells, surface area increases linearly with radius, whereas volume increases in proportion to the radius squared. Consequently, larger guard cells will require more time, all else being equal, to achieve a given change in turgor pressure and, hence, stomatal pore aperture. A plant with large guard cells can partially compensate for this geometric constraint by increasing the activity and/or density of ion and solute transporters in the guard cell plasma membrane (Homann & Thiel, 2002; Lawson & Blatt, 2014; Raven, 2014). However, compensation may be limited by the available membrane area and/or energetic costs of maintaining high transporter densities (Vico *et al*., 2011). Empirical support for the hypothesis that smaller guard cells are faster is mixed and has primarily come from step changes in light (Drake *et al*., 2013; Elliott-Kingston *et al*., 2016; McAusland *et al*., 2016; Kardiman & Ræbild, 2018; Haworth *et al*., 2023; Yoshiyama *et al*., 2024; Brench *et al*., 2026; Tang *et al*., 2026). Meta-analysis of dozens of species suggests that stomatal size alone is weakly related to kinetics and modulated by guard cell shape (Woning *et al*., 2026). The relationship between guard cell size and stomatal response kinetics to leaf water potential (Aasamaa *et al*., 2001) and VPD (Kübarsepp *et al*., 2020; Qie *et al*., 2024) is less well studied, but similarly equivocal.

One factor that might complicate the relationship between size and speed is aperture. Stomatal aperture can vary from near 0 when guard cell turgor pressure is low and asymptotically approach *g*_max_ as guard cell turgor pressure approaches ∞ (Figure 1d). The relationship between guard cell turgor pressure and stomatal aperture is nonlinear (Franks & Farquhar, 2001) as cell walls become less elastic at greater turgor (Franks *et al*., 2001). The nonlinear relationship between guard cell turgor and pore aperture implies that if the rate of turgor change is constant 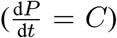, the rate of change in aperture, and hence stomatal conductance, will vary depending on the initial aperture (*a*_0_) relative to the maximum aperture (*a*_max_). Hence, 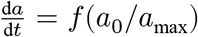. When initial aperture is high, *g*_sw_ will respond more slowly than when initial aperture is low. Since we generally do not observe individual stomatal aperture, relating this idea to leaf-level *g*_sw_ requires scaling by stomatal density, biophysical constants, and morphological constants (e.g. Sack & Buckley, 2016). We will use the term “fraction of anatomical maximum stomatal conductance”, symbolized as *f*_gmax_ (this is denoted as *g*_ratio_ by Xie *et al*. (2022)). This value should be proportional to the average aperture divided by its maximum aperture for any arbitrary stomatal density. Leaves operating at *f*_gmax_ closer to unity will, all else being equal, be slower to respond than leaves operating with a *f*_gmax_ close to zero, independent of stomatal density. Franks *et al*. (2012) hypothesized that selection on stomatal density would maintain typical operational stomatal conductance (*g*_op_) relative to *g*_max_ at a sufficiently low value that *g*_sw_ would be sensitive to relatively small changes in guard cell turgor pressure. Consistent with this hypothesis, the *g*_op_:*g*_max_ ratio is often near 0.25 (McElwain *et al*., 2016; Murray *et al*., 2020), well below *f*_gmax_ = 1, and in a range where *g*_sw_ would be responsive to small changes in guard cell turgor pressure. There is likely variation within and among species and growth forms in the *g*_op_:*g*_max_ ratio (Xie *et al*., 2022; Ochoa *et al*., 2024).

**Figure 1:**
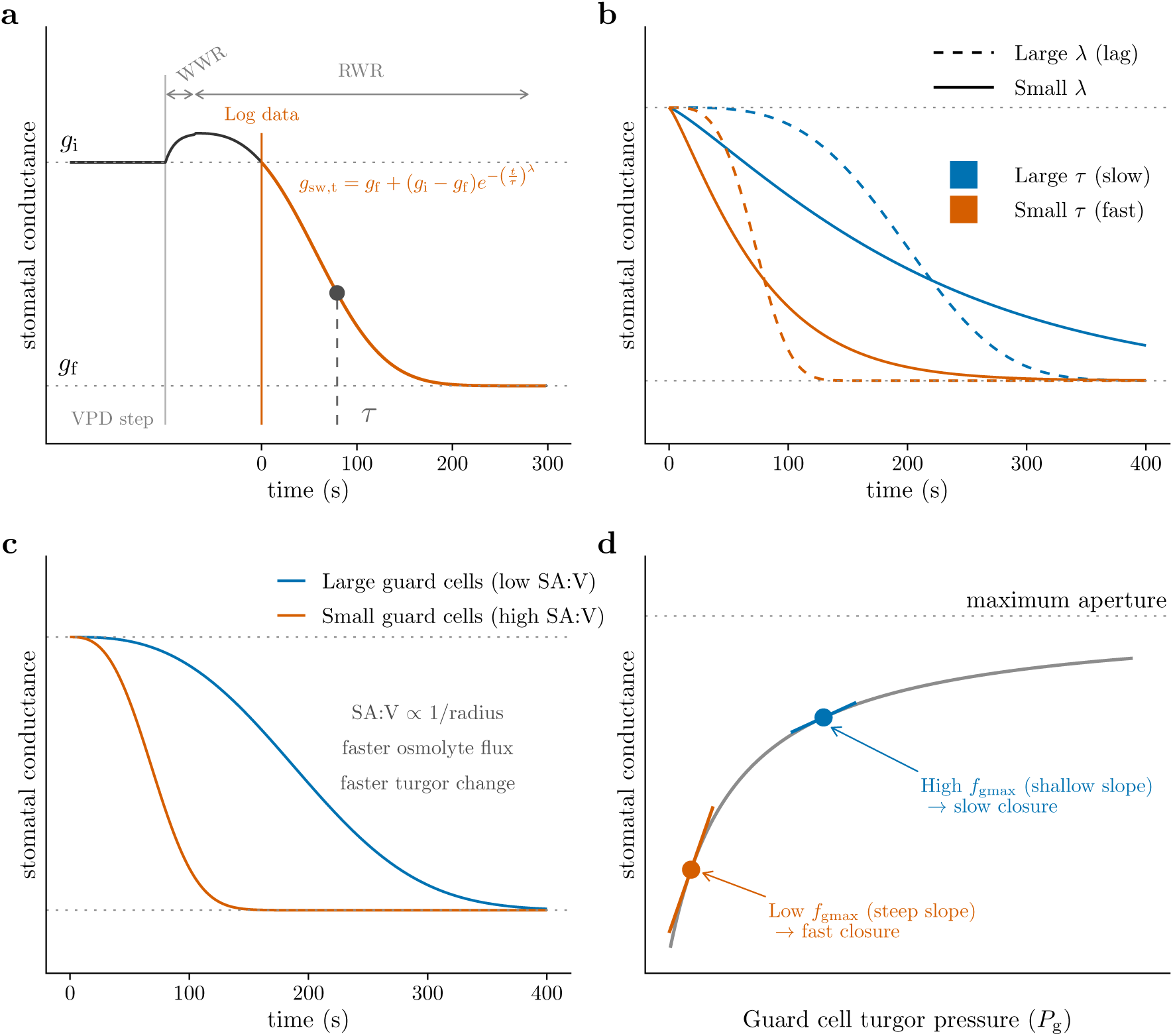
Conceptual overview of the two hypotheses linking stomatal anatomy to closure kinetics. **a.** Stomatal conductance (*g*_sw_) in response to a step increase in vapor pressure deficit (VPD) typically exhibits a transient wrong-way response (WWR), in which *g*_sw_ briefly rises, followed by the right-way response (RWR), in which *g*_sw_ declines to a lower steady state. The RWR is well described by a Weibull function (dashed line; Equation 1) with initial conductance *g*_i_, final conductance *g*_f_, and kinetic parameters *τ* and *λ*. **b**. The time constant *τ* sets the overall speed of closure, and the lag time *λ* determines whether the response rate is roughly constant (*λ*≈ 1; solid) or accelerates over time (*λ*> 1; dashed). **c**. Smaller guard cells have a greater surface area to volume ratio (SA:V; schematic cross-sections shown at right) and are predicted to exchange osmolytes more quickly across the plasma membrane, yielding a faster change in guard cell turgor pressure and faster closure. **d**. The relationship between guard cell turgor pressure (*P*_*g*_) and *g*_sw_ is nonlinear and saturating because guard cell walls become less elastic at higher turgor (Franks & Farquhar, 2001; Franks *et al*., 2001). Consequently, the same change in turgor pressure produces a larger change in *g*_sw_ when stomata are at low *f*_gmax_ (orange; steep tangent slope) than at high *f*_gmax_ (blue; shallow tangent slope), predicting slower closure when *f*_gmax_ is high.

Putting these two hypotheses together, we predict that both guard cell size and *f*_gmax_ will influence stomatal kinetics, but possibly at different scales of biological organization. Guard cell size tends to vary less than stomatal density or aperture at many biological scales. For example, on a mature leaf, guard cell length is nearly constant even as cell volume and pore aperture change with turgor (Willmer & Fricker, 1996; Franks *et al*., 2001). Compared to stomatal density, guard cell size can be less variable (Lammertsma *et al*., 2011; Bucher *et al*., 2016) and evolve more slowly between species (Muir *et al*., 2023). If there is relatively little plastic or genetic variation among individuals within a species, then most of the variance in stomatal kinetics within species cannot be explained by variation in guard cell size. In contrast, *f*_gmax_ can vary immensely within and among individuals because stomatal aperture responds dynamically over the day and in response to environmental variation. Because guard cell size and aperture typically vary at different levels of biological organization, we predict that guard cell size likely explains more variation in stomatal kinetics among species, whereas *f*_gmax_ will explain more variation within species. Within species variation can arise from either genetic differences among individuals or environmental plasticity. We will also test if adaxial stomata respond faster than abaxial stomata (Haworth *et al*., 2018; Wall *et al*., 2022; Woning *et al*., 2026) and, if so, whether this difference can be explained by variation in guard cell size and/or *f*_gmax_ between surfaces.

One challenge in determining whether traits covary at higher levels of organization is that statistical procedures to account for phylogenetic nonindependence can obscure ecologically relevant covariation caused by natural selection (Westoby *et al*., 1995; Uyeda *et al*., 2018). For example, in a meta-analysis of stomatal kinetic responses to light, the explanatory power of traits like guard cell size and stomatal ratio declined in phylogenetic compared to nonphylogenetic regression (Woning *et al*., 2026). This finding may be caused in large part because grasses have small stomata, high adaxial:abaxial stomatal ratio, and rapid stomatal kinetics. Are these associations a coincidence caused by shared ancestry or ecologically important covariation that helps explain how grasses adapt to their environment? Recent advances in phylogenetic comparative methods use hierarchical, multiresponse phylogenetic models (Muir *et al*., 2023; Westoby *et al*., 2023; Halliwell *et al*., 2025) to tease apart “conservative trait correlations” that act at higher levels of organization (e.g. among species) from trait correlations that explain variation at lower levels of organization (e.g. among individuals within species and plastic responses). We use this approach to test whether guard cell size and *f*_gmax_ explain variation in stomatal kinetics at different levels of biological organization. Throughout, we use guard cell length (*l*_gc_) as a proxy for size (Sack & Buckley, 2016).

The four main questions we address in this study are:

1. Are VPD kinetics adequately characterized by the same functional form in Equation 1 as light responses?
2. Are *l*_gc_ and *f*_gmax_ positively associated with *τ* ?
3. At what biological levels of organization do *l*_gc_ and *f*_gmax_ vary and influence *τ* ?
4. Are abaxial-only stomatal responses slower than whole leaf stomatal responses?

As noted above, we are opportunistically analyzing data from an experiment designed to answer different questions. Before analyzing the kinetic data, we planned to test whether 1) guard cell size would influence *τ* and 2) that *τ* would be greater on the abaxial-only leaves based on a recent meta-analysis of stomatal responses to light (Woning *et al*., 2026). We made the decision to test for an association between *f*_gmax_ and *τ* after exploratory analyses revealed a potential association. Adding hypotheses after some results have been analyzed, also known as “Hypothesizing After Results Known” (HARKing), can lead to reports of spurious associations that are not repeatable in later studies (Kerr, 1998; Smaldino & McElreath, 2016). Therefore, our conclusions about the effect of *f*_gmax_ on stomatal kinetics should be interpreted with caution until they are replicated.

## Materials and Methods

### Populations and phylogeny

We grew 29 populations of wild tomato (Table S1), including representatives of all described species of *Solanum* sect. *Lycopersicon* and sect. *Lycopersicoides* (Peralta *et al*., 2008), as described in Muir *et al*. (2025). These populations were selected because they encompass the breadth of climatic variation in the wild tomato clade (Pease *et al*., 2016). Due to constraints on growth space and time, we spread out measurements over 61.1 weeks. Replicates within population were evenly spread out over this period to prevent confounding of temporal variation in growth conditions with variation among populations. For phylogenetic comparative analyses, we used the RAxML whole-transcriptome concatenated phylogeny based on 2,745 100-kb genomic windows from Pease *et al*. (2016). Two of our populations were not in this tree. We used *S. galapagense* accession LA1044 in place of LA3909. *S. pennellii* accession LA0750 was added as sister to LA0716. The node separating LA0750 from LA0716 was placed halfway along the branch to the next deepest node.

### Plant growth conditions and light treatments

A thorough description of plant growth conditions can be found in Muir *et al*. (2025), therefore we summarize the most important information here. Seeds provided by the Tomato Genetics Resource Center germinated on moist paper in plastic boxes after soaking for 30-60 minutes in a 50% (volume per volume) solution of household bleach and water, followed by a thorough rinse. We transferred seedlings to cell-pack flats containing Pro-Mix BX potting mix (Premier Tech, Rivière-du-Loup, Quebec, Canada) once cotyledons fully emerged, typically within 1-2 weeks of sowing. We grew seeds and seedlings for both sun and shade treatments under the same environmental conditions (12:12 h, 24.3:21.7 °C, 49.6:58.4 RH day:night cycle). LED light provided PPFD = 267 µmol m^−2^ s^−1^ (Fluence RAZRx, Austin, Texas, USA) during the germination and seedling stages.

Immediately prior to starting growth light intensity treatments (sun and shade), we transplanted seedlings to 3.78 L plastic pots containing 60% Pro-Mix BX potting mix, 20% coral sand (Pro-Pak, Honolulu, Hawai‘i, USA), and 20% cinders (Niu Nursery, Honolulu, Hawai‘i, USA) with slow release NPK fertilizer following manufacturer instructions (Osmocote Smart-Release Plant Food Flower & Vegetable, The Scotts Company, Marysville, Ohio, USA). Percentage composition is on a volume basis. We watered to field capacity three times per week to prevent drought stress. Seedlings were randomly assigned in alternating order within population to the sun or shade treatment during transplanting. The average daytime PPFD was 761 µmol m^−2^ s^−1^ and 115 µmol m^−2^ s^−1^ for sun and shade treatments, respectively.

### Humidity response curves

We selected a fully expanded, unshaded leaf at least six leaves above the cotyledons during early vegetative growth. This typically meant that plants had grown in light treatments for ≈ 4 weeks, ensuring they had time to sense and respond developmentally to the light intensity of the treatment rather than the seedling conditions (Schoch *et al*., 1980).

We measured stomatal conductance over time after a step change in humidity at a constant leaf temperature, which we refer to this as a humidity-response curve. We measured humidity-response curves using a portable infrared gas analyzer (LI-6800PF, LI-COR Biosciences, Lincoln, Nebraska, USA). During gas exchange measurements, light-acclimated plants were placed under LEDs dimmed to match their light treatment. We collected four humidity-response curves per leaf, an amphistomatous (untreated) curve and a pseudohypostomatous (treated) curve at high light-intensity (PPFD = 2000 µmol m^−2^ s^−1^; 97.8:2.24 red:blue) and low light-intensity (PPFD = 150 µmol m^−2^ s^−1^; 87.0:13.0 red:blue). We always measured high light-intensity curves first because photosynthetic downregulation is faster than upregulation in these species. As a consequence, measurement light intensity is confounded with measurement order in our design: low light-intensity curves were always measured after high light-intensity curves on the same leaf, following a low-humidity closure event, so we cannot fully separate an effect of light intensity *per se* from order, recovery state, or prior VPD exposure (see Discussion). Pseudohypostomatous leaves are the same as the amphistomatous leaves except that gas exchange through the upper (adaxial) surface is blocked by a neutral density plastic (propafilm). For brevity, we hereafter refer to these as “pseudohypo” and “amphi” leaf types. Comparing amphi and pseudohypo leaf types enabled us to test whether stomatal kinetics are different for ab- and adaxial stomata on the same leaf. To compensate for reduced transmission, we increased incident PPFD for pseudohypo leaves by a factor 1/0.91, the inverse of the measured transmissivity of the propafilm. We also set the stomatal conductance ratio, for purposes of calculating boundary layer conductance, to 0 for pseudohypo leaves following manufacturer directions. To control for order effects, we alternated between starting with amphi or pseudohypo measurements. We made measurements over two days. On the first day, we measured high and low light-intensity curves for either amphi or pseudohypo leaves; on the second day, we measured high and low light-intensity curves on the other leaf type at approximately the same time of day.

In all humidity-response curves, we acclimated the focal leaf ambient CO_2_ (*C*_a_ = 415 µmol mol^−1^), *T*_leaf_ = 25.0 °C, high light (PPFD = 2000 µmol m^−2^ s^−1^), and high relative humidity (RH = 70%; VPD ≈ 0.82 kPa) until *g*_sw_ reached its maximum. After that, we scrubbed water vapor from the incoming air using silica desiccant as quickly as possible, resulting in a water vapor concentration in the incoming air to be on average 0.36 mmol mol^−1^. This treatment induced rapid stomatal closure, but it did not standardize chamber air humidity because leaf transpiration introduced water vapor. Consequently, the RH was systematically higher at high light intensity, in sun plants, and amphi leaf types (Table S2) because the *g*_sw_ was typically greater in these conditions. In all treatment conditions, the final RH (5.37% − 13.8%) and VPD (2.72 kPa −3 kPa) elicited rapid stomatal response and is comparable to treatments used in similar experiments (Merilo *et al*., 2014; Kübarsepp *et al*., 2020; Hsu *et al*., 2021; Qie *et al*., 2024), but was not intended to mimic natural environmental variability. Because the realized VPD stimulus varied among treatments, we checked whether this could confound treatment, *f*_gmax_ components, and *l*_gc_ effects on stomatal closure kinetics using curve-level VPD covariates (Pichaco *et al*., 2024). We found that our conclusions were robust to this potential confound (Notes S1; Figs. S1–S4). Following the transient WWR, we started logging data once *g*_sw_ declined back to its pre-step value (*g*_i_) and continued logging data until *g*_sw_ reached its nadir (Figure 1a). We then acclimated the leaf to low light (PPFD = 150 µmol m^−2^ s^−1^) and RH = 70% before inducing stomatal closure with low RH and logging data again, on average 48.9 minutes after the start of the high-light curve.

Prior to analysis, we removed unreliable and unusable data points, removed points not at instrument steady state, removed the hysteretic portion of each curve at low *g*_sw_, removed outliers within each curve, removed replicates with no overlap between amphi and pseudohypo curves from the same leaf, and thinned redundant data points within each curve. The rationale and procedure for each cleaning step is described in Muir *et al*. (2025). Additionally, we excluded stomatal kinetic parameter estimates from five curves where *τ* was estimated to be greater than 1097 s, far exceeding the typical estimates (see Results). After these steps, the total number of humidity response curves analyzed is given in Table S3. The median number of curves per population per treatment was 9 and the median number of points per curve was 26. We extracted the posterior distribution for *τ* and *λ* from each fitted curve to calculate the parameter estimate (median) and uncertainty (standard deviation) for subsequent analyses.

### Stomatal anatomical traits *l*_gc_ and *f*_gmax_

We estimated the stomatal density and *l*_gc_ on ab- and adaxial surfaces from all leaves used for humidity response curves using standard methods described fully in Muir *et al*. (2025). We excluded three individuals with extreme outlying guard cell length values. Preliminary analysis of variation in guard cell length among growth treatments, leaf types, and accessions identified these outliers based on a residual value greater than 0.45, which visually appeared as a clear break in the residual distribution. When the leaflet used for the LI-6800 humidity-response curve did not have a matching leaflet-level anatomical measurement, we used the first available anatomical measurement from a different leaflet on the same leaf as a proxy; this fallback was used for 9 of 561 individual-by-leaf-type combinations (1.6%). For each surface we calculated the anatomical maximum stomatal conductance (*g*_max_) at a reference leaf temperature of 25 °C following Sack & Buckley (2016) as:

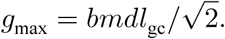

The biophysical and morphological constants *b* and *m* are:

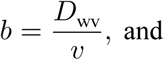

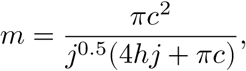

where *D*_wv_ is the diffusion coefficient of water vapor in air and *v* is by the kinematic viscosity of dry air. We assumed *D*_wv_ = 2.49 × 10^−5^ m^2^ s^−1^ and *v* = 2.24 × 10^−2^ m^3^ mol^−1^ (Monteith & Unsworth, 2013). For kidney-shaped guard cells like those in wild tomatoes, *c* = *h* = *j* = 0.5. To estimate *f*_gmax_, we divided the initial *g*_sw_ estimated from the parameter *g*_i_ in Equation 1 from each humidity response curve by the anatomical *g*_max_ for that leaf. The estimated *g*_i_ was very close to the maximum *g*_sw_ observed near the beginning of each curve (Figure S5), suggesting that the estimated *g*_i_ was a good proxy for stomatal conductance at the time of the step change in humidity. Because *g*_i_ is estimated from the same nonlinear fit as *τ* and *λ*, we used a null simulation to confirm that this does not, by itself, induce a spurious *f*_gmax_-*τ* association (Notes S2). We used the anatomical *g*_max_ for the surface(s) measured in each curve to calculate *f*_gmax_. For amphi curves, we used the sum of adaxial and abaxial *g*_max_; for pseudohypo curves, we used only the abaxial *g*_max_. Although *g*_max_ calculated this way is a theoretical upper bound based on stomatal geometry rather than a directly measured quantity, it corresponds well to empirically measured maximum *g*_sw_ in experimentally controlled *Arabidopsis* (Dow *et al*., 2014).

### Estimating kinetic parameters

We estimated *τ* and *λ* for each response curve by fitting Equation 1 to a time course of *g*_sw_ values using Bayesian nonlinear regression. All models were fit using Bayesian HMC sampling in the probabilistic programming language *Stan* (Stan Development Team, 2026) using the *R* package **brms** version 2.23.0 (Bürkner, 2017). We used *CmdStan* version 2.38.0 and **cmdstanr** version 0.9.0 (Gabry *et al*., 2025) to interface with *R* version 4.6.1 (R Core Team, 2026). We sampled 4,000 post-warmup draws from the posterior distribution across four chains. We adjusted the number of sampling iterations and thinning interval for each curve so that the Gelman-Rubin convergence statistic 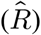 (Gelman & Rubin, 1992) was less than 1.05 and the bulk effective sample size (ESS) was greater than 400 for all model parameters, and there were fewer than 10 divergent transitions. To answer Question 1, we assessed model fit using Bayesian *R*^2^ (Gelman *et al*., 2019) implemented in **brms**.

### Estimating effects of stomatal anatomy on kinetic parameters

We first used model selection to determine whether anatomy (*l*_gc_ and *f*_gmax_) affected kinetic parameters (*λ* and *τ*; Question 2) and at what level of biological organization (Question 3). To infer the effect of *f*_gmax_, we modeled effects of component traits log(*g*_i_) and log(*g*_max_) directly because log (*f*_gmax_) = log (*g*_i_) − log (*g*_max_). If *f*_gmax_ *per se* affects kinetics then both log(*g*_i_) and log(*g*_max_) should be in the selected model. Alternatively, *g*_i_ or *g*_max_ could effect kinetic parameters independent of *f*_gmax_. From the selected model, we used parameter estimates and 95% confidence intervals to test the predictions of our hypotheses. We interpreted nonzero fixed effects of *l*_gc_, *g*_i_, and *g*_max_ on *⋋* and *τ* as evidence that these anatomical traits influence variation in stomatal kinetics among individuals. We interpreted nonzero phylogenetic correlations among traits as evidence that these traits covary among populations. We predicted that *l*_gc_ and *g*_i_ should be positively associated with *λ* and *τ*, whereas the effect of *g*_max_ should be negative. Finally, we used path analysis to test whether treatment-associated plasticity in *τ* was statistically consistent with an indirect path through *l*_gc_ and/or *f*_gmax_ components. Table 1 summarizes the statistical criteria for determining whether and at what level of biological organization stomatal anatomical traits affected kinetic parameters. The subsections below describe specific procedures for deciding whether those are met.

**Table 1:** Criteria used to determine whether and at what level of biological organization stomatal anatomical traits affected kinetic parameters.

| Level of organization | Criteria supporting an association between stomatal anatomy and kinetics |
| --- | --- |
| Among individuals | • Anatomical parameters as fixed effects improve model fit (lower LOOIC)<br>• Anatomical effects on kinetics are in the predicted direction<br>• Confidence intervals of anatomical effects do not overlap zero |
| Plasticity among individuals | • Plasticity in anatomy is statistically consistent with an indirect path underlying plasticity in stomatal kinetics |
| Among populations | • Confidence intervals of phylogenetic correlation among traits do not overlap zero<br>• Phylogenetic covariance is in the predicted direction |

### Bayesian multiresponse phylogenetic models

Multiresponse models can disentangle whether traits are associated among populations, individuals, and/or treatments that induce plasticity. The response variables in our models are *τ*, *⋋, l*_gc_, *g*_i_, and *g*_max_. All models included a subset of explanatory variables that were integral to the experimental design or necessary to account for statistical nonindependence. As explanatory variables, we included fixed effects of manipulative treatments: growth light intensity (sun and shade); measurement light intensity (high and low); and leaf type (amphi and pseudohypo). All models also included population as both a phylogenetically structured random effect and a non-phylogenetic random effect following Halliwell *et al*. (2025). Prior to fitting, the phylogenetic (co)variance matrix was rescaled as a correlation matrix (De Villemereuil & Nakagawa, 2014). For traits estimated from multiple curves within the same leaf (*λ, τ*, *g*_i_) we included a random effect of individual to account for repeated measures. We accounted for uncertainty in *τ* and *λ* using the standard deviation of the posterior distribution of each estimate (see ‘Humidity response curves’ for details). We modeled residual (co)variance, a combination of among-individual (co)variance and measurement error, as a multivariate Student *t*-distribution, which is more robust to extreme values than the multivariate normal distribution (Gelman *et al*., 2014). To account for heteroskedasticity, bounded measurements, and to linearize relationships all response variables were log-transformed prior to analysis. All models were fit with *Stan* on three chains using the same methods and convergence criteria described above for fitting Equation 1 to the humidity response curves.

### Phylogenetic correlation among traits

We inferred that traits covary among populations if the 95% confidence intervals of the phylogenetic correlation between traits did not overlap zero and the correlation was in the predicted direction. For example, a positive phylogenetic correlation between *l*_gc_ and *τ* would be consistent with the prediction that species with larger guard cells have slower stomatal kinetics. Phylogenetic covariance specifically indicates long-term evolutionary covariation among traits because our model simultaneously accounts for trait associations among individuals, shared plastic responses, and residual covariance. To test for possible causation, we used the inferred phylogenetic trait covariance matrix to estimate partial correlations between anatomical and kinetic variables (Halliwell *et al*., 2025). We interpret a significant partial correlation between variables as a plausible causal hypothesis.

### Model selection and parameter estimation of individual-level effects

To infer whether individual-level variation in *l*_gc_ and *f*_gmax_ influence kinetic parameters, we used the leave-one-out cross-validation information criterion (LOOIC) to compare the fit of models using the *R* package **loo** version 2.10.1 (Vehtari *et al*., 2017). We compared models with all permutations of *l*_gc_, *g*_i_, and *g*_max_ influencing *⋋* and *τ*. We considered all models within two ΔLOOIC standard errors of the minimum LOOIC (Table S4) to be in the set of plausible models to generate posterior predictions for hypothesis testing. Within each plausible model, we assessed whether a fixed effect was significantly different than zero based on the 95% confidence intervals. Among the set of plausible models, we selected the model with the highest proportion of significant focal parameters for further analysis. Focal parameters are the fixed and phylogenetic effects of *l*_gc_, *g*_i_, and *g*_max_ on *⋋* and *τ*. This approach aids interpretability because it favors simpler models that exclude additional nonsignificant parameters. Because this choice is somewhat arbitrary and not based on rigorous statistical principles, we discuss alternative interpretations based on the other plausible models. We report Pareto 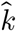, PSIS effective sample size, and Monte Carlo standard error diagnostics for the plausible models in Table S5. Nearly every curve had good Pareto 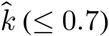, but the minimum PSIS effective sample size per curve was sometimes low (as low as 94 draws for some models), so the PSIS-LOO estimates for a small number of individual curves, and by extension the model-level Monte Carlo standard error, should be interpreted with some caution.

### Path analysis to test for plastic effects

We used path analysis (Pearl, 2022) to test whether treatment-associated plasticity in *τ* was statistically consistent with an indirect path through plasticity in stomatal traits, in response to measurement light intensity, growth light intensity, and curve-type treatments. We focused on *g*_i_ because it was the only individual-level trait significantly associated with *τ* in the selected model (see Results). We do not treat this as a randomized-treatment causal mediation analysis because *g*_i_ is not randomized, is estimated from the same curve as *τ*, and could share unmeasured confounders (e.g., hydraulic status, measurement order, realized VPD trajectory) with *τ*, violating the sequential ignorability assumption required for a causal interpretation. We assessed the plausibility of this assumption directly with a sensitivity analysis (Notes S3). We inferred the direct effect of treatments on *τ* from model coefficients. We estimated the indirect effect of treatments as the product of the effect of treatment on *g*_i_ and the effect of that trait on *τ* (Imai *et al*., 2010). We determined significant direct and indirect effects based on whether the 95% confidence intervals of the effects were different than zero. We calculated effect sizes as the point estimate (median of the posterior) divided by the standard error (standard deviation of the posterior).

### Variance decomposition and phylogenetic heritability

One explanation for why traits have effects at different levels of organization is that variance in those traits is concentrated at those different levels of organization. To evaluate this, we decomposed the variance in response variables (*λ, τ*, *l*_gc_, *g*_i_, *g*_max_) into phylogenetic, population (nonphylogenetic), and individual levels. The phylogenetic variance component quantifies trait evolution between populations congruent with the phylogenetic relationships, whereas the population (nonphylogenetic) component quantifies variance among populations independent of phylogenetic relationships (Lynch, 1991; De Villemereuil & Nakagawa, 2014). We estimated these components from the associated random-effect standard deviations for each response variable. The among-individual variance components quantify variance among individuals within the same population and treatment. We estimated the among-individual component as the sum of the individual random-effect and residual standard deviations. It was not possible to estimate individual random-effects in *l*_gc_ or *g*_max_ with our study design because we only measured average guard cell length and *g*_max_ once per leaf. The among-individual component is biased upwards because it also includes measurement error, which cannot be separately estimated with our experimental design. The phylogenetic heritability is equivalent to the phylogenetic variance component and is estimated following Lynch (1991) and De Villemereuil & Nakagawa (2014) as:

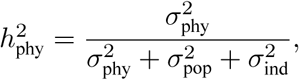

where 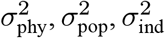 are the phylogenetic, population (nonphylogenetic), and among-individual variance components, respectively. Because phylogenetic heritability was estimated after accounting for fixed treatment effects, estimates obtained in controlled environments likely overestimate phylogenetic heritability in natural environments, where plasticity is a larger component of trait variation.

### Effect of adaxial stomata on kinetics

If adaxial stomata respond faster than abaxial stomata (Question 4), we predicted that *τ* should be greater in pseudohypo leaves compared to amphi. We evaluated this prediction by estimating the fixed effect of leaf type (amphi vs pseudohypo) on *τ* in the multiresponse model. We did not estimate adaxial-only kinetics because it was not part of the original experimental design (Muir *et al*., 2025).

## Results

### The time constant *τ* explains most variation in stomatal closure kinetics

Stomatal conductance (*g*_sw_) decreased rapidly in response to a step change in humidity. The model in Equation 1 fit the data extremely well (Figure S6). The mean and minimum Bayesian *R*^2^ across all curves was 0.9994 and 0.9803, respectively. Consider a typical leaf in the experiment characterized by the median time constant (*τ*) and lag time (*λ*) among wild tomato populations in all treatments. After humidity decreased, the transient WWR elapsed, and *g*_sw_ returned to the initial value *g*_i_, it took on average 128 s for *g*_sw_ to decrease half-way (*t*_50_) from its initial to final (*g*_f_) steady state value. In terms of the kinetic model parameters, most variation among leaves was due to difference in the time constant rather than the lag time. In the previous example, increasing *λ* from its median to its maximum estimated value among populations increased the *t*_50_ by only 7.1 s. In contrast, increasing *τ* from its median to maximum value increased *t*_50_ by 86 s. Hence, we primarily focus on treatments and anatomical parameters affecting *τ*.

### Variation in guard cell length and *f*_gmax_ components

Guard cell length (*l*_gc_) differed among populations and varied to a lesser extent between growth light intensity treatments and leaf surfaces. In contrast, stomatal conductance as a fraction of maximum conductance (*f*_gmax_) was most strongly determined by measurement light intensity. Because *g*_max_ is unaltered by measurement light intensity, this treatment effect was entirely caused by changes in *g*_i_. Excluding treatment effects, phylogenetically structured differences among populations (i.e., phylogenetic heritability) accounted for 61.9% [95% CI: 18.9% to 82.8%] of the variance in *l*_gc_ (Figure 2, Table S6). There was also developmental plasticity in *l*_gc_. On average, *l*_gc_ increased by 12.1% [95% CI: 11.1% to 13.1%] in the sun treatment (Figure 3a; Table S7). The average *l*_gc_ of pseudohypo leaves was shorter by −5.8% [95% CI: −6.6% to −5.0%] compared to amphi leaves (Figure 3a; Table S7). This was not because of plasticity, as we measured amphi and pseudohypo curves on the same leaf. Rather, this was a composition effect. Adaxial stomata are larger than abaxial stomata in these species (Figure S7), so the average *l*_gc_ in pseudohypo leaves was lower than in amphi leaves because the former only included abaxial stomata.

**Figure 2:**
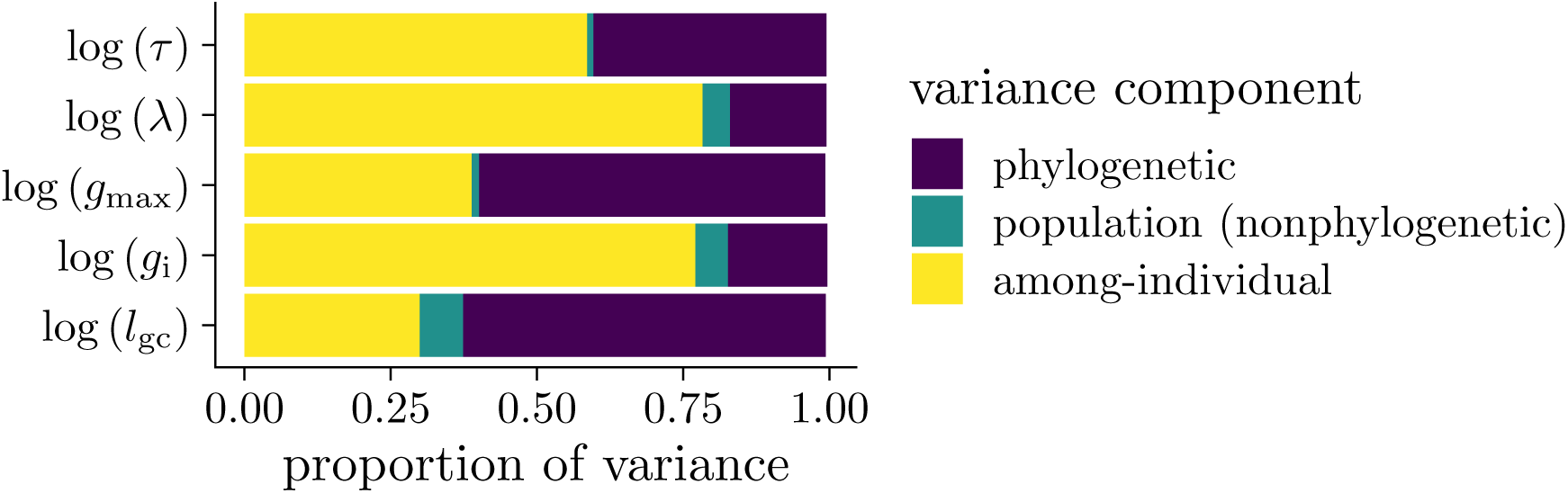
Traits vary at different levels of biological organization. A large fraction of the variation in guard cell length (*l*_gc_), anatomical maximum stomatal conductance (*g*_max_), and the time constant *τ* is phylogenetically structured among populations. In contrast, most of the variance in initial stomatal conductance (*g*_i_) and the lag time (*⋋*) is among individuals. Nonphylogenetic variation among populations was low for all traits.

**Figure 3:**
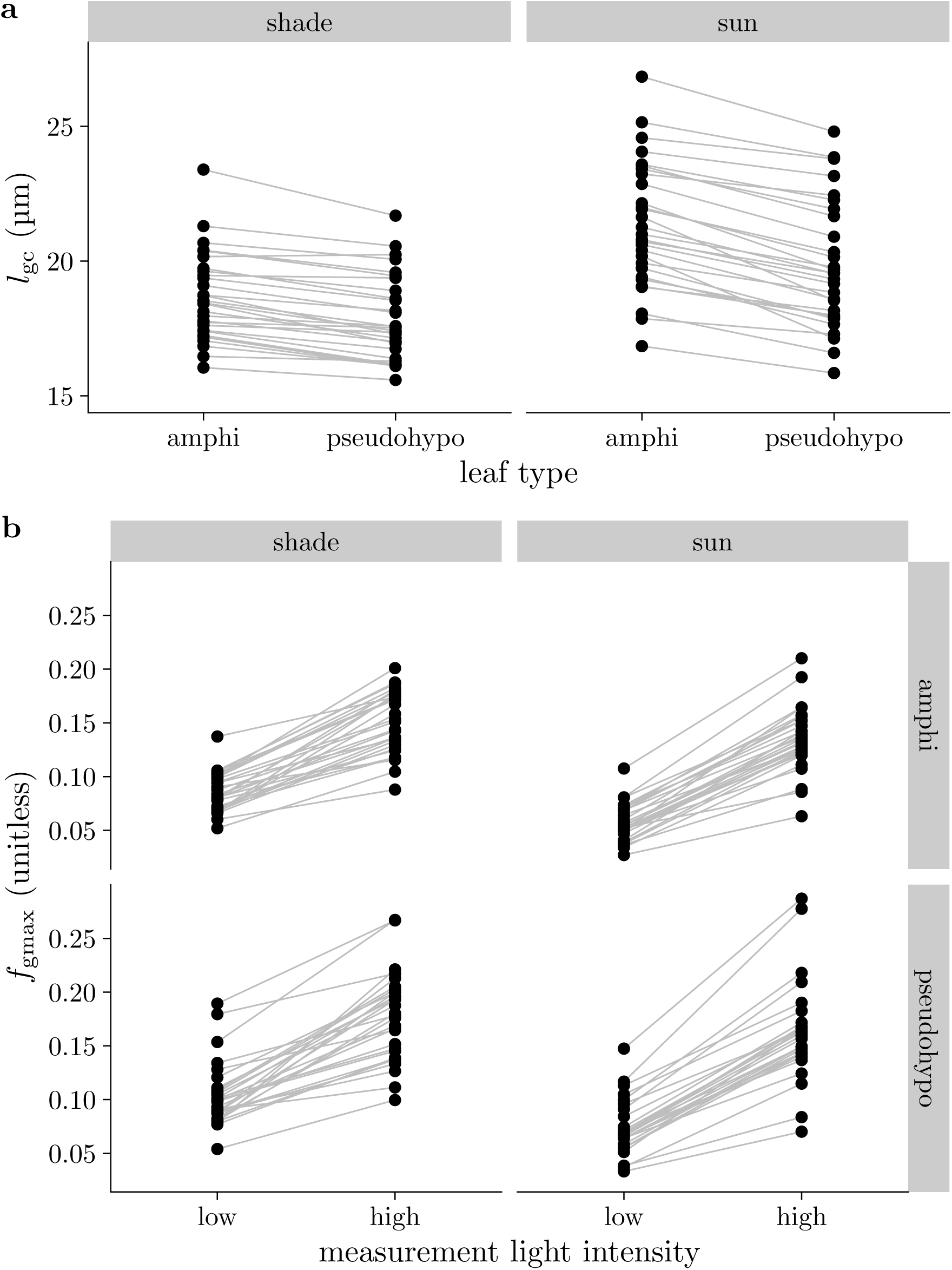
Guard cell length (*l*_gc_) and stomatal conductance as a fraction of maximum conductance (*f*_gmax_) vary among populations and treatments. In both panels, black points are mean trait value per population within a particular treatment. Gray lines connect values within populations across treatments. (a) Most of the variation in *l*_gc_ is among populations, but developmental plasticity to growth light intensity tends to increase *l*_gc_ in sun (right facet) compared to shade (left facet) grown plants. There is no plasticity to leaf type (amphi vs. pseudohypo), but the average *l*_gc_ is slightly lower in pseudohypo leaves because adaxial stomata tended to be larger than abaxial stomata on the same leaf. (b) Measurement light intensity has the most direct effect on *f*_gmax_ because initial stomatal conductance (*g*_i_) increases under high light intensity. To a lesser extent, *f*_gmax_ varies among populations, is slightly lower in sun-grown plants, and slightly higher in pseudohypo leaf types.

The phylogenetic heritability of *f*_gmax_ components was low (*g*_i_) to moderate (*g*_max_), accounting for 16.9% [95% CI: 1.1% to 37.9%] and 59.1% [95% CI: 34.4% to 75.5%] of the variance, respectively (Figure 2, Table S6). High measurement light intensity increased *g*_i_ by an average of 113.2% [95% CI: 107.7% to 118.5%] (Figure 3b; Table S7). Sun leaves had on average 41.6% [95% CI: 34.4% to 49.5%] greater *g*_max_ than shade-grown plants (Figure 3b; Table S7). The pseudohypo leaf type had on average −19.5% [95% CI: −21.5% to −17.3%] lower *g*_i_ than untreated, amphistomatous leaves (Figure 3b; Table S7). This is presumably because a large fraction of stomata were blocked by the pseudohypo treatment and increased aperture of the abaxial stomata did not fully compensate. This observation differs from wheat, where *g*_sw_ on one surface was not affected by blocking gas exchange in the other (Wall *et al*., 2022).

### Variation in stomatal closure kinetic parameters

Estimates of stomatal closure kinetic parameters for each curve are in Table S8 and estimates for each population are in Table S9. The time constant *τ* exhibited moderate phylogenetic heritability, accounting for 39.8% [95% CI: 18.7% to 60.4%] of the variance, whereas *λ* varied more within populations (Figure 2, Table S6). Some light treatments also affected kinetic parameters. The *τ* in sun plants was −10.7% [95% CI: −15.1% to −6.06%] lower than shade plants after accounting for other explanatory variables (Figure 4, Table S7). Within the same leaf, increasing the measurement light intensity from 150 µmol m^−2^ s^−1^ to 2000 µmol m^−2^ s^−1^ significantly increased *τ* by 8.6% [95% CI: 1.4% to 16.4%] (Figure 4, Table S7). The lag time *λ* responded to growth, but not measurement light intensity after accounting for other explanatory variables (Figure 4, Table S7). In sun plants, *λ* was on average 3.43% [95% CI: 1.4% to 5.67%] greater than in shade plants. High measurement light had little effect on *λ*, with an estimated change of −1.55% [95% CI: −4.35% to 1.39%].

**Figure 4:**
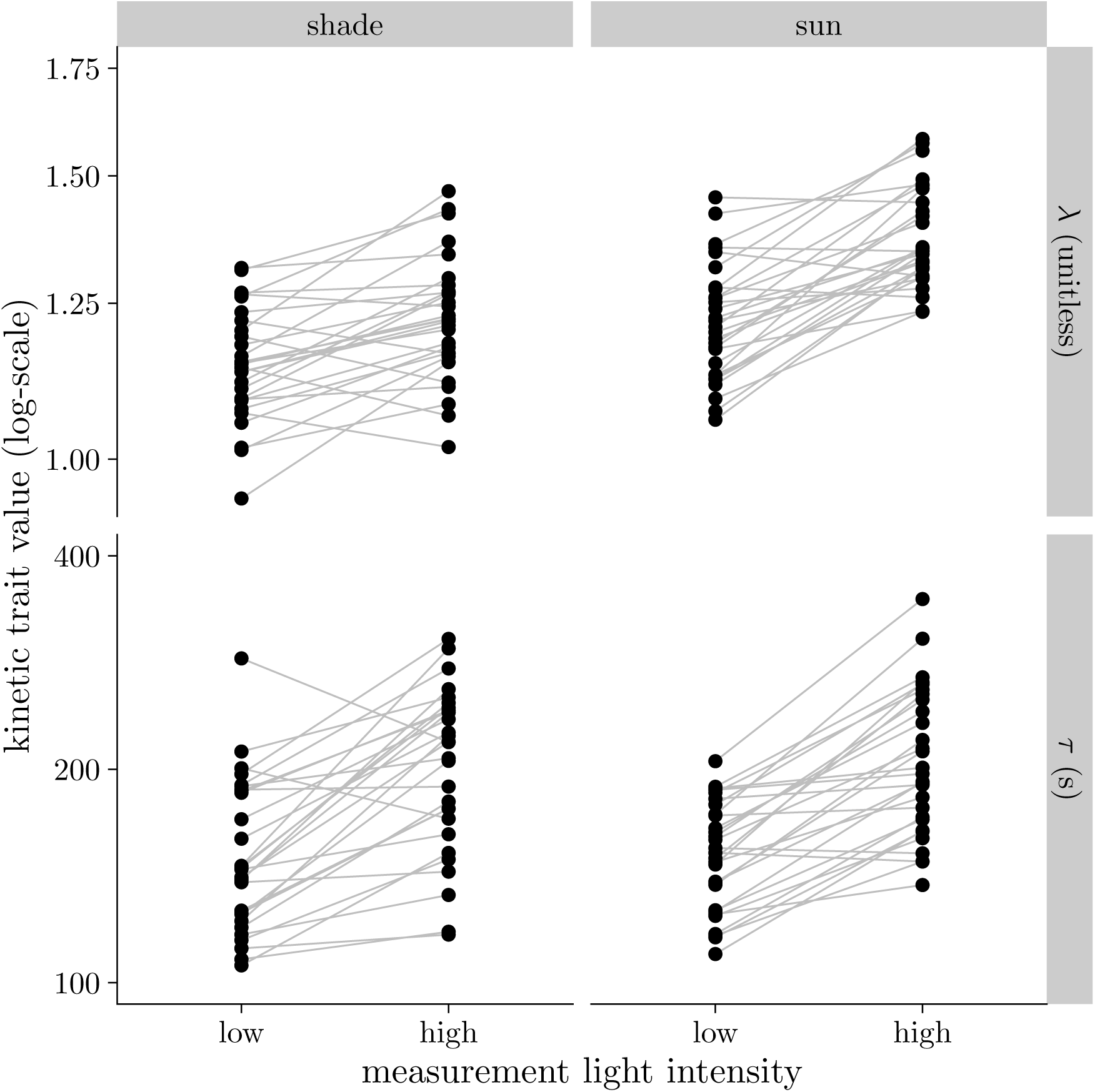
Stomatal kinetics parameters *⋋* (lag time; top facets) and *τ* (time constant; lower facets) vary among wild tomato populations. Stomatal conductance decreases faster (lower *τ*) in response to a step change in vapor pressure deficit (VPD) in leaves when measured under low light intensity. The pattern was consistent for both shade (left facets) and sun (right facets) grown plants. The pattern for *⋋* was qualitatively similar to that for *τ*. Each point is the average parameter value for one accession in that treatment combination. The growth and measurement light intensity treatments are described in the Materials and Methods section.

### Guard cell length and *g*_i_ influence stomatal kinetics

Guard cell length and *g*_i_, a component of *f*_gmax_, affected stomatal closure kinetics in the directions predicted by our hypotheses, but they operated at different levels of biological organization. Among the models we tested, six were plausible in that their predictive accuracy was similar based on ΔLOOIC (Table S4). All models in the plausible set indicated among-individual level effects of *g*_i_ on *τ* and *⋋*, as we discuss in more detail below. Models that included both individual and phylogenetic effects of *l*_gc_ had similar LOOIC values but the selected model retained only phylogenetic effects because our selection criteria favored models that excluded nonsignificant parameters (Table S4). In models that included both individual and phylogenetic effects, neither the phylogenetic correlation nor the fixed effect regression coefficient were significantly different than zero (Figure S8; Figure S9). These parameters are highly collinear in the posterior (Figure S10) indicating difficulty in estimating both terms simultaneously and “smearing” out their confidence intervals (McElreath, 2020). Hence, the model with only phylogenetic effects is easier to interpret. This model is also consistent with the high phylogenetic heritability in both *l*_gc_ and *τ* (Figure 2). Visually, we observed positive relationships among population mean trait values (Figure 5) but little relationship between *l*_gc_ and *τ* within populations (results not shown). When comparing many models simultaneously, differences in LOOIC performance can occur by chance (McLatchie & Vehtari, 2024), so judgement calls among models with similar performance can be necessary. The preponderance of evidence indicates that *l*_gc_ explains variation primarily among rather than within our populations. The results in the main text are based on estimates from the selected model. Individual-level fixed effects (Figure S8) and phylogenetic correlations (Figure S9) of *l*_gc_, *g*_i_, and *g*_max_ on *τ* and *⋋* were qualitatively consistent across all plausible models.

**Figure 5:**
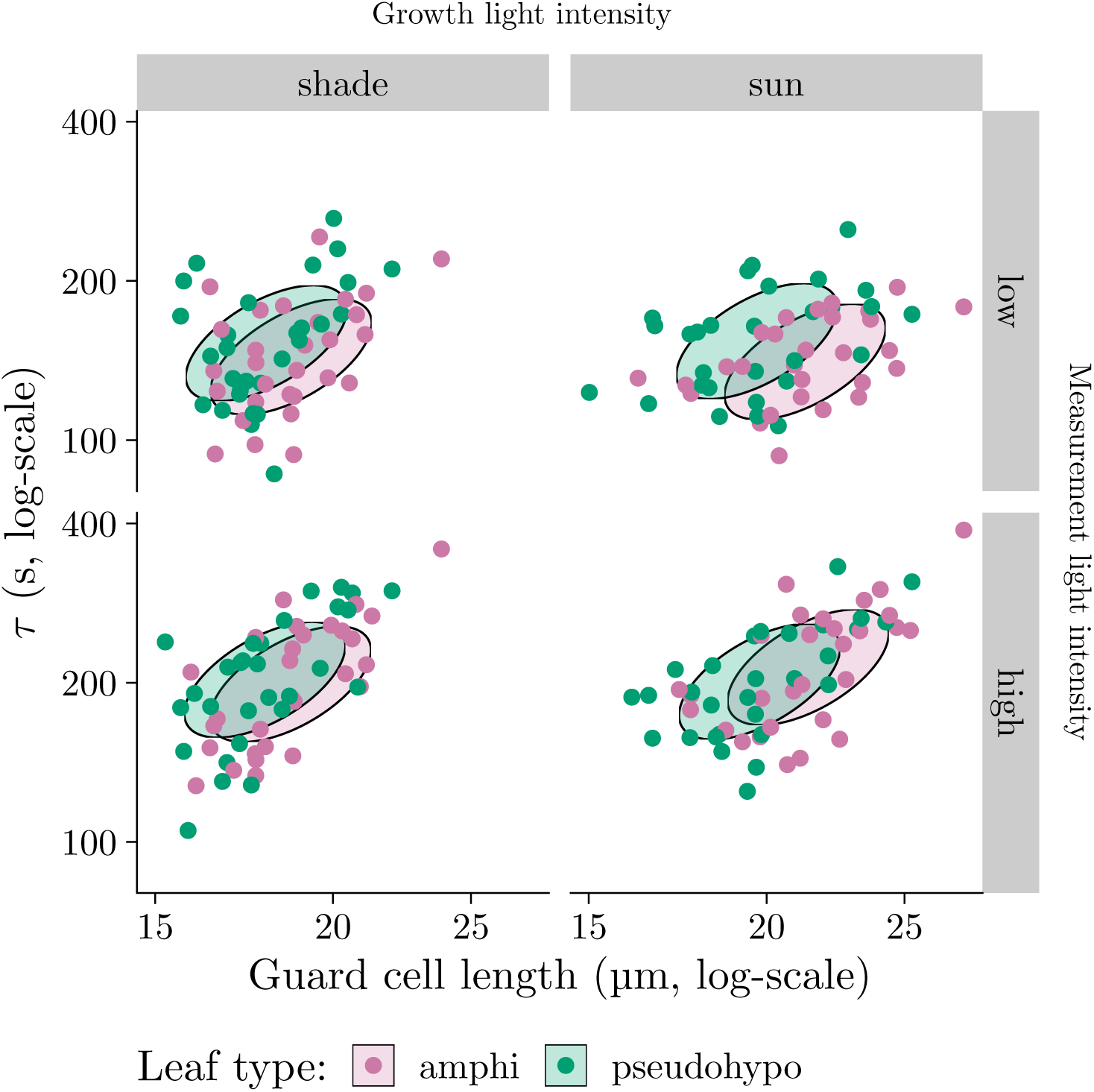
Phylogenetic covariance between guard cell length (*l*_gc_, *x*-axis) and the stomatal closure time constant (*τ*, *y*-axis) indicates that populations with larger stomata tend to close more slowly in response to humidity. The overall pattern was consistent across growth light intensity treatments (left and right facets), measurement light intensity treatments (top and bottom facets), and leaf types (point colors). The direction and magnitude of the ellipses are such that they should encompass 95% of the phylogenetically structured (co)variance in *l*_gc_ and *τ* on log-transformed scales. The location of ellipses are set to the median trait values in each treatment combination for visual comparison. Each point is the average parameter value for one population in that treatment combination. The growth and measurement light intensity treatments are described in the Materials and Methods section.

Leaves with greater *g*_i_ closed more slowly than those with lower *g_i_* (Figure 6; Table S7). For example, increasing *g*_i_ from 0.2 to 0.4 mol m^−2^ s^−1^ resulted in a 25.1% [95% CI: 17.6% to 32.6%] increase in *τ* for a typical leaf in the reference treatment set (shade, low, amphi). We found little evidence that *g*_max_ affected *τ* (Figure S11). Leaves with greater *g*_i_ also had a significantly greater lag time, *λ* (Figure S12), but there was little effect of *g*_max_ (Figure S13).

**Figure 6:**
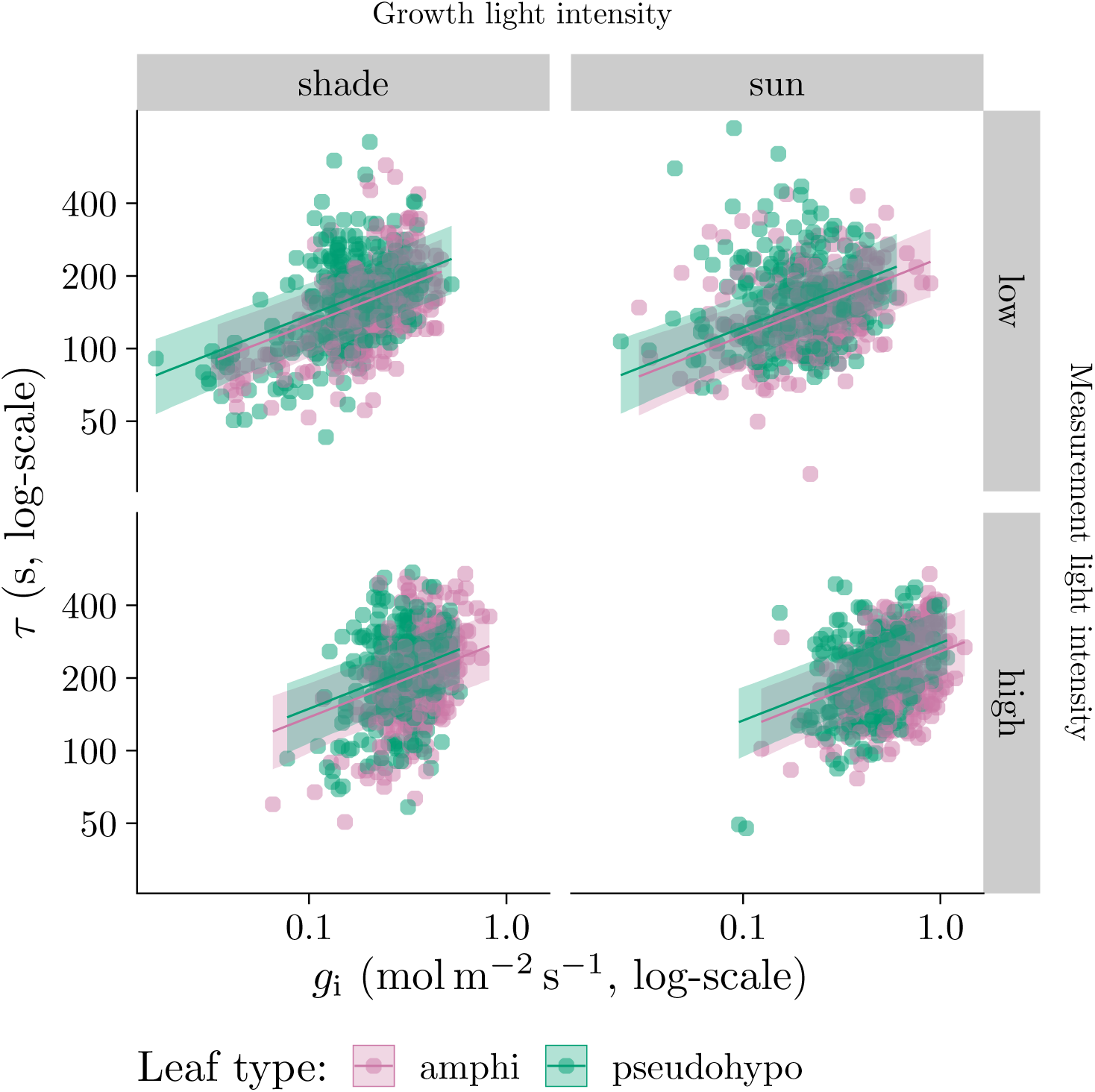
Individual-level variation in initial stomatal conductance (*g*_i_) influences stomatal closure kinetics. As *g*_i_ (*x*-axis) increases, the stomatal closure time constant (*τ*, *y*-axis) increases. The overall pattern was consistent across grow light intensity treatments (left and right facets), measurement light intensity treatments (top and bottom facets), and leaf types (point colors). Lines are estimated using linear regression along with 95% confidence ribbons. The growth and measurement light intensity treatments are described in the Materials and Methods section.

Populations that tended to have greater *l*_gc_ also tended to have greater *τ* as evidenced by a positive phylogenetic correlation (Table S10, Figure 5). However, we found little evidence that variation in *l*_gc_ explained *τ* or *⋋* within populations. There were no other significant phylogenetic correlations (Table S10). The only marginally significant partial correlation was between *l*_gc_ and *τ* (*ρ* = 0.71 [95% CI: −0.17 to 0.98]), indicating a plausible causal effect of *l*_gc_ on *τ* among populations.

### Plasticity in *τ* is an indirect effect of plasticity in *g*_i_

We imposed three treatments that influenced stomatal kinetics: growth light intensity, measurement light intensity, and leaf type. All three treatments directly affected *τ* and had indirect effects mediated by *g*_i_ (Figure 7). We did not estimate effects mediated by *l*_gc_ because the selected model did not retain direct effects of *l*_gc_ on *τ* (Table S4). The indirect effect of greater *g*_i_ in sun leaves increased *τ* by 11.78% [95% CI: 8.27% to 15.99%], counteracting the direct negative effect. The direct effect of the pseudohypo treatment increased *τ* by 8.60% [95% CI: 5.62% to 11.56%], but the indirect effect mediated by lower *g*_i_ changed *τ* by −6.72% [95% CI: −8.64% to −4.83%]. In contrast, direct and indirect effects of high measurement light intensity increased *τ*. The direct effect increased *τ* by 8.56% [95% CI: 1.40% to 16.40%] and the indirect effect mediated by *g*_i_ increased *τ* by 27.62% [95% CI: 19.18% to 36.24%].

**Figure 7:**
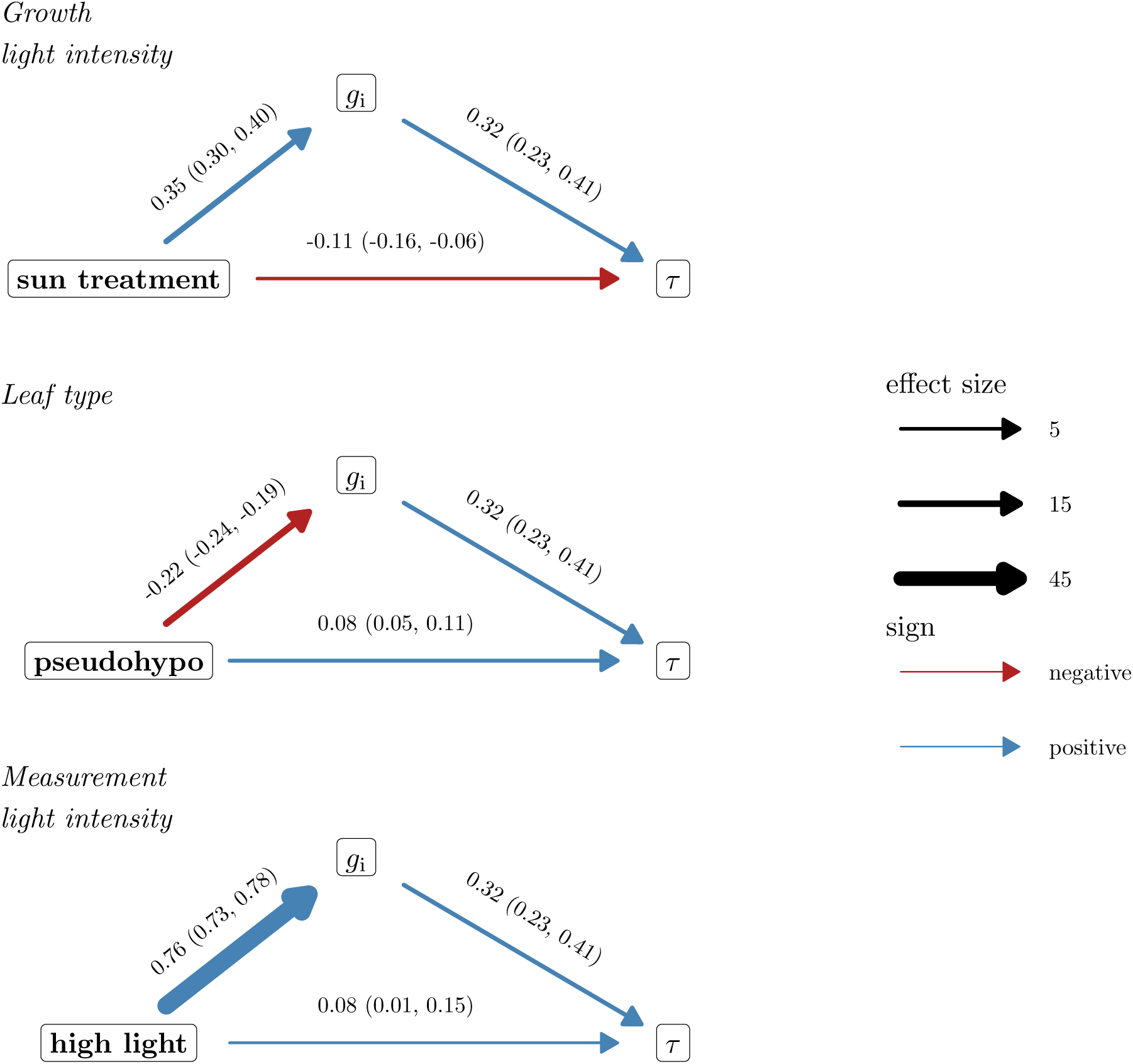
Treatment-associated plasticity in the time constant *τ* is statistically consistent with an indirect path through initial stomatal conductance (*g*_i_). The three treatments are growth light intensity (top facet; shade vs. sun), leaf type (middle facet; amphi vs. pseudohypo), and measurement light intensity (bottom facet; low vs. high). Horizontal arrows at the bottom of each graph indicate the direct effect of the treatment level on *τ*; slanted arrows leading to or from *g*_i_ indicate indirect effects of the treatment on *τ* through *g*_i_. Line thickness is proportional to the effect size and line color indicates whether the effects are negative (red) or positive (blue). Parameter estimates of the effect and 95% confidence intervals are above each line. The selected model did not retain direct effects of *l*_gc_ on *τ* and hence this variable could not be included in this analysis. This path analysis assumes sequential ignorability (no unmeasured confounder of the *g*_i_-*τ* relationship); see Notes S3 for a sensitivity analysis of this assumption.

### Adaxial stomata close faster than abaxial stomata, but the effect is masked by plasticity in *g*_i_

Although a simple comparison of *τ* between paired amphi and pseudohypo leaf types shows little net difference (results not shown), the selected model reveals a significant direct effect of leaf type on *τ* once *g*_i_ is accounted for: pseudohypo leaves, which have only abaxial stomata, closed more slowly than amphi leaves (Figure 7; Table S7). This direct effect is consistent with the hypothesis that adaxial stomata close faster than abaxial stomata. However, pseudohypo leaves also had lower *g*_i_ and lower *g*_i_ is itself associated with faster closure (Figure 6). This indirect path through *g*_i_ counteracts most of the direct effect (Figure 7), which is why the raw comparison of paired leaves does not reveal much difference in kinetics between leaf types even though adaxial stomata appear to close faster once *g*_i_ is controlled for.

## Discussion

The ability for stomata to adjust to fluctuating environmental conditions likely evolved in early land plants because of selection to optimize water use relative to carbon gain (Raven, 2002). Hydroactive control of stomatal aperture in angiosperms increases flexibility to modulate carbon gain and water loss depending on leaf water status (Buckley, 2019), perhaps enabling a competitive advantage in environments where light and evaporative demand fluctuate at high frequency (McAdam & Brodribb, 2012, 2015; Pichaco *et al*., 2026). We now understand the steady-state response of stomatal conductance (*g*_sw_) to global increases in vapor pressure deficit (VPD) better but there is greater uncertainty surrounding kinetic responses to VPD (Buckley *et al*., 2023). Here we considered two nonmutually exclusive mechanistic hypotheses for how stomatal anatomy influences stomatal closure kinetics. First, smaller stomata may close faster because of greater surface area to volume ratio (Drake *et al*., 2013). Second, guard cells under higher turgor pressure respond more slowly because of reduced cell wall elasticity (Franks *et al*., 2012). Based on these hypotheses, we predicted that leaves with smaller guard cells and operating further from their anatomical maximum *g*_sw_ (i.e., lower *f*_gmax_) would have faster stomatal closure kinetics (lower *τ*). The parameter *f*_gmax_ is here defined as the initial stomatal conductance (*g*_i_) divided by *g*_max_. Analysis of 2,125 time courses of stomatal closure from 29 populations in multiple environments demonstrates that both guard cell size and *g*_i_, a component of *f*_gmax_, influence kinetics at different levels of biological organization. This new understanding suggests unexplored explanations for the relationships between stomatal size, density, speed, and operational *g*_sw_ (*g*_op_) among angiosperms.

The kinetics of stomatal closure in response to VPD closely match response to light, suggesting similar underlying mechanisms. The lag-time model proposed by Woning *et al*. (2026) for light responses also describes humidity responses extremely well following the passive wrong-way response (WWR). The fitted models explained on average 99.94% of the variation in *g*_sw_ during closure (Figure S6). The formal mathematical similarity of light and humidity response might indicate that certain factors, such as guard cell size, influence stomatal kinetics responses in general. Alternatively, disparate underlying processes may be described by the same macroscale formalism. For example, humidity responses may be controlled by ABA accumulation or loss (McAdam *et al*., 2016), whereas *C*_i_-independent red-light responses can be influenced by accumulation or loss of photosynthetic products (Matthews *et al*., 2020). These disparate processes may both have similar kinetic properties, but would not necessarily imply that VPD and light responses are strongly correlated. Future research comparing the correlation between light, VPD, CO_2_, and other stomatal kinetic responses can determine the extent to which generic or specific factors determine stomatal opening and closing rates (Creese *et al*., 2014; Kübarsepp *et al*., 2020).

The positive association between *l*_gc_ and *τ* (Figure 5) is consistent with the hypothesis that smaller cells respond faster because they have a greater surface area to volume ratio for osmolyte flux (Franks & Farquhar, 2007; Drake *et al*., 2013; Lawson & Vialet-Chabrand, 2019; Woning *et al*., 2026). Among several of the same *Solanum* species, larger guard cells are also associated with slower opening kinetics in response to light (Yoshiyama *et al*., 2024), suggesting a general mechanism. Specifically, this association implies that the maximum rate of stomatal closure is limited by membrane surface area. We do not have the data to evaluate the hypothesis mechanistically, but this should be tested in the future. In addition to guard cell size, it will be useful to test whether the ratio of guard cell to adjacent epidermal size affects the strength of the WWR, which may be necessary to induce more rapid stomatal closure (Buckley *et al*., 2003; Pichaco *et al*., 2025). An alternative explanation is that the correlation between *l*_gc_ and *τ* is not causal. The fact that *l*_gc_ did not explain much of the variation in *τ* among individuals within populations suggests that *l*_gc_ may not directly affect *τ*. However, the high phylogenetic heritability and, hence, low variability in *l*_gc_ within populations may also explain why we were unable to detect a within-population effect of *l*_gc_ on *τ*.

A potential consequence of greater stomatal aperture is that the pore closes more slowly when the environment changes. Much of the variation in *τ* among individuals within populations was explained by variation in *g*_i_ (Figure 6). This association is not simply an artifact of the magnitude of the difference between *g*_i_ and *g*_f_, because *τ* captures the proportional rate of change, not the absolute change, in *g*_sw_. This factor was positively associated with *τ* among individuals within light treatments, and treatment-associated plasticity in *τ* was statistically consistent with an indirect path through *g*_i_ (Figure 7). The rate of stomatal opening in response to light is associated with *g*_op_ in a sample of rainforest trees (Kardiman & Ræbild, 2018), which may indicate this is a more general phenomenon.

The association between *g*_i_ and *τ* could be explained by the hypothesis that guard cell wall elasticity declines at greater turgor pressure (Franks *et al*., 2012), like a rubber band that is stretched under tension. However, if this were the case, we would also expect a negative association between *g*_max_ and *τ* because leaves with greater *g*_max_ could have the same *g*_i_ with a smaller stomatal aperture. However, this effect was weakly supported (Figure S11, Figure S13). The selected model did not include an individual-level effect of *g*_max_ on kinetic parameters. Although the second-best model included *g*_max_ and the effect was in the predicted direction (Table S4), the effect was not significantly different than 0 (Figure S8). The weak effect of *g*_max_ compared to *g*_i_ casts doubt on whether a nonlinear relationship between turgor pressure and pore aperture is the best explanation. An alternative, albeit not mutually exclusive explanation, is a nonlinear relationship between pore aperture and *g*_sw_ that weakens the association between these variables at high *g*_sw_. Airspace resistance within the substomatal cavity and stomatal interference are plausible mechanisms that would have this effect (Ting & Loomis, 1965; Kaiser, 2009; Lehmann & Or, 2015), although it is not clear that they are sufficient to produce the observed relationship between *g*_i_ and *τ* in wild tomatoes. Our study is limited by the fact that we did not directly measure guard cell turgor or stomatal aperture, but future studies should use newly developed methods (Brodersen *et al*., 2025) to directly evaluate the association between these traits.

The impact of *g*_i_ was most pronounced when comparing measurement light intensity treatments. Leaves acclimated to high light intensity had on average 113% greater *g_i_* than the same leaves at low light intensity in the reference treatment (shade, amphi). As a consequence, stomata closed more slowly in leaves acclimated to high light intensity, with a 27.6% greater time constant on average in the reference treatment, based on parameter estimates from the selected model (Table S7). We did not assess whether *g*_i_ also affects stomatal opening and this should be measured to further test this hypothesis. The implication is that natural selection should favor *g*_op_ values that not only optimize instantaneous carbon gain and water loss in the present environment, but also carbon gain and water loss integrated through time in a fluctuating environment. We predict that rapid, temporally-uncorrelated fluctuating environments will favor leaves with greater *g*_op_ to slow kinetic responses and prevent overreacting to transitory changes that are likely to revert; slower, temporally-correlated changes will favor lower *g*_op_ to enable close tracking of the environment.

The comparison between measurement light intensity treatments should be interpreted cautiously, however, because it was confounded with measurement order: high light curves always preceded low light curves on the same leaf, following a prior low-humidity closure event (see Materials and Methods). We cannot rule out that some of the apparent effect of measurement light intensity on *g*_i_ and *τ*, and the corresponding indirect path through *g*_i_ in our mediation analysis (Figure 7), reflects order, incomplete recovery, or prior VPD exposure rather than measurement light intensity itself. This concern applies specifically to the measurement light comparison and not to growth light intensity or leaf type, which were randomized or alternated, respectively.

The direct effect of leaf type on *τ* indicates that adaxial stomata close faster than abaxial stomata after accounting for differences in *g*_i_. This result is consistent with the broader association between greater adaxial stomatal density and faster stomatal closure across land plants (Haworth *et al*., 2018; Woning *et al*., 2026). One plausible reason adaxial stomata may have evolved to close intrinsically faster is because adaxial stomata directly overlie the photosynthetically active palisade mesophyll and failure to close quickly could rapidly desiccate this tissue and impair photosynthesis (Tezara *et al*., 1999; Buckley *et al*., 2017a). Hypostomatous leaves may experience less dehydration risk because the palisade is hydraulically buffered from changing VPD when stomata are confined to the abaxial surface (Buckley *et al*., 2017a). If traits such as guard cell size that influence kinetics act similarly on both surfaces, this shared basis, together with the intrinsic speed difference between surfaces documented here, could jointly explain why leaves and species with a greater proportion of adaxial stomata tend to have faster overall stomatal kinetics.

Future research is needed to determine how stomatal kinetic responses measured in controlled conditions impact performance in nature. Our experimental protocol induced stomatal closure by abruptly switching incoming air to near-zero humidity, a perturbation far more severe than the gradual or partial VPD fluctuations leaves experience in the field. This approach is well suited for estimating the maximum rate of stomatal closure and comparing it among individuals and treatments, but it need not follow that the same anatomical and physiological factors that govern *τ* in our experiment also govern kinetic variation under more naturalistic conditions, where closure VPD may fluctuate more gradually. Future studies should also control the VPD stimulus throughout the entire response curve, rather than allowing it to vary as transpiration itself alters chamber humidity, as it did here (Table S2). It will also be important to consider whether the substomatal airspace remains saturated during the humidity step, because *g*_sw_ will be underestimated if the airspace is subsaturated (Cernusak *et al*., 2018; Buckley & Sack, 2019; Wong *et al*., 2022; Rockwell *et al*., 2022). We hypothesize that ecological conditions which favor close tracking of fluctuating environments will select for leaves with traits like smaller guard cell size and low *g*_i_ that enable rapid kinetics.

## Acknowledgements

Comments from two anonymous reviewers greatly improved the manuscript. Sam McKlin and Tom Buckley helped with protocol development. Justin Alter, Max Gatlin, Joana Kim, Jenna Matsuyama, Brandon Najarian, Dachuan Wang, and Kai Yasuda contributed to data collection. We used Copilot, ChatGPT, and Posit Assistant (Claude) to assist with coding and proofreading text.

## Competing interests

None declared.

## Funding

US National Science Foundation OIA-1929167 (C.D.M.)

## Author contributions

Conceptualization: C.D.M.; Data acquisition: C.D.M., W.S.L.; Data analysis and visualization: C.D.M.; Writing – original draft: C.D.M.; Writing – review & editing: C.D.M., W.S.L.

## Data availability

All data and code are available on GitHub (https://github.com/cdmuir/solanum-kinetics). Data are archived on Dryad (https://doi.org/10.5061/dryad.f7m0cfzcv); code is archived on Zenodo (https://doi.org/10.5281/zenodo.22802259).

## Notes

https://github.com/cdmuir/solanum-kinetics/

### Competing Interest Statement

The authors have declared no competing interest.

### Summary of Updates

We changed 36.8% to 63.2% in the Introduction to fix an error. The main text incorrectly stated that “Sun leaves had on average 41.6%[95% CI: 34.4% to 49.5%] greater *g*max than shade-grown plants.” This was caused by copy-and-paste error in the code that we have now corrected. We clarified that the 27.6% figure was specifically the indirect effect of high measurement light intensity on τ mediated by *g*i, and we now also report the combined total effect (direct + indirect), which is 38.6% [95% CI: 35.3% to 41.7%]. We note in the text that direct and indirect percentage effects combine additively on the log scale, not the percentage scale, which is why they do not simply sum to the total. We clarified that "much" or "a large fraction" of the variation in in guard cell length was phylogenetic, rather than saying "most" of the variation. We replaced references to more or less "open" with references to *g*i specifically. We fixed the null simulation code to analyze the correlation between log (*g*i) and log (τ) for consistency throughout the manuscript.

