## Supporting Information for "Guard cell size and initial conductance influence stomatal closure kinetics"

### Supporting tables

Table S1: Accession information of *Solanum* populations used in this study. The species name, accession number, collection latitude, longitude, and elevation. TGRC: Tomato Genetics Resource Center; mas: meters above sea level.

| Species | TGRC accession | Latitude | Longitude | Elevation (mas) |
|---|---|---|---|---|
| *S. arcanum* | LA2172 | -6.008 | -78.858 | 662 |
| *S. cheesmaniae* | LA0429 | -0.644 | -90.329 | 800 |
| *S. cheesmaniae* | LA3124 | -0.804 | -90.042 | 1 |
| *S. chilense* | LA1782 | -15.267 | -74.633 | 1000 |
| *S. chilense* | LA4117A | -22.907 | -67.941 | 3540 |
| *S. chmielewskii* | LA1028 | -13.883 | -73.017 | 3000 |
| *S. chmielewskii* | LA1316 | -13.400 | -73.906 | 2920 |
| *S. corneliomulleri* | LA0107 | -13.117 | -76.383 | 60 |
| *S. corneliomulleri* | LA0444 | -13.433 | -76.133 | 100 |
| *S. galapagense* | LA0436 | -0.953 | -90.978 | 40 |
| *S. galapagense* | LA1044 | -0.284 | -90.548 | 0 |
| *S. habrochaites* | LA0407 | -2.181 | -79.884 | 70 |
| *S. habrochaites* | LA1777 | -9.550 | -77.700 | 3216 |
| *S. huaylasense* | LA1358 | -9.533 | -77.967 | 750 |
| *S. huaylasense* | LA1360 | -9.546 | -77.929 | 1490 |
| *S. huaylasense* | LA1364 | -10.133 | -77.383 | 2920 |
| *S. lycopersicoides* | LA2951 | -19.317 | -69.450 | 2200 |
| *S. lycopersicoides* | LA4126 | -19.287 | -69.396 | 3120 |
| *S. neorickii* | LA1322 | -13.483 | -72.442 | 2380 |
| *S. neorickii* | LA2133 | -3.400 | -79.183 | 1980 |
| *S. pennellii* | LA0716 | -16.225 | -73.617 | 50 |
| *S. pennellii* | LA0750 | -14.775 | -75.034 | 550 |
| *S. pennellii* | LA3778 | -14.775 | -75.034 | 616 |
| *S. peruvianum* | LA2744 | -18.550 | -70.150 | 400 |
| *S. peruvianum* | LA2964 | -18.028 | -70.835 | 75 |
| *S. pimpinellifolium* | LA1269 | -11.483 | -77.075 | 400 |
| *S. pimpinellifolium* | LA1589 | -8.433 | -78.817 | 30 |
| *S. pimpinellifolium* | LA2933 | -1.442 | -80.562 | 375 |
| *S. sitiens* | LA4116 | -22.159 | -68.782 | 2960 |

Table S2: Relative humidity (RH, %) and leaf vapor pressure deficit (VPD, kPa) during stomatal kinetics measurements systematically differed because of differences in stomatal conductance. Stomatal conductance tended to be greatest in sun-grown plants, measured at high light intensity, and untreated (amphistomatous). For each treatment combination, we report the mean across replicate curves of the RH and VPD at the start (Initial) and end (Final) of the fitted interval, and of the median RH and VPD across time points within the fitted interval (Median).

| | | | RH | | | VPD | | |
|---|---|---|---|---|---|---|---|---|
| Growth light intensity | Measurement light intensity | Leaf type | Initial | Median | Final | Initial | Median | Final |
| shade | low | pseudohypo | 12 | 7.62 | 5.37 | 2.78 | 2.92 | 3.00 |
| shade | low | amphi | 13.8 | 8.25 | 5.68 | 2.72 | 2.91 | 2.99 |
| shade | high | pseudohypo | 18.4 | 13.3 | 9.98 | 2.58 | 2.74 | 2.85 |
| shade | high | amphi | 21.2 | 14.7 | 10.6 | 2.48 | 2.69 | 2.83 |
| sun | low | pseudohypo | 13.8 | 9.63 | 7.39 | 2.72 | 2.85 | 2.93 |
| sun | low | amphi | 16.7 | 11.1 | 8.23 | 2.62 | 2.81 | 2.91 |
| sun | high | pseudohypo | 24.3 | 16.8 | 11.6 | 2.36 | 2.62 | 2.79 |
| sun | high | amphi | 29.8 | 20.4 | 13.8 | 2.17 | 2.50 | 2.72 |

Table S3: Total number of humidity response curves analyzed in this study broken down by treatments.

| | Measurement light intensity | | | | |
|---|---|---|---|---|---|
| | Low | | High | | |
| Growth light intensity | amphi | pseudohypo | amphi | pseudohypo | Total |
| shade | 275 | 275 | 270 | 266 | 1,086 |
| sun | 245 | 246 | 274 | 274 | 1,039 |
| Total | 520 | 521 | 544 | 540 | 2,125 |

Table S4: Model comparison results. Models with a ΔLOOIC less than two standard errors (SE) from the minimum LOOIC are considered to have substantial support and are used for posterior predictions to test hypotheses about the effects of guard cell length ($l_{gc}$) and the components ($g_i$, $g_{max}$) of stomatal conductance as a fraction of anatomical maximum stomatal conductance ($f_{gmax}$) on stomatal kinetic parameter $\lambda$ (lag time) and $\tau$ (time constant). Unsupported models are in gray italic font. The model selected for subsequent inference is in bold. LOOIC stands for leave-one-out cross-validation information criterion.

| | | $\lambda$ | | | $\tau$ | | |
|---|---|---|---|---|---|---|---|
| ΔLOOIC | SE | $l_{gc}$ | $g_i$ | $g_{max}$ | $l_{gc}$ | $g_i$ | $g_{max}$ |
| **0.00** | **0.00** | | ✓ | | | ✓ | |
| 3.58 | 2.95 | ✓ | ✓ | ✓ | ✓ | ✓ | ✓ |
| 4.80 | 2.96 | | ✓ | | ✓ | ✓ | |
| 4.91 | 2.92 | | ✓ | | | ✓ | ✓ |
| 4.92 | 2.90 | ✓ | ✓ | | | ✓ | |
| 5.20 | 2.72 | | ✓ | ✓ | | ✓ | |
| *5.91* | *2.91* | ✓ | ✓ | ✓ | | ✓ | ✓ |
| *6.06* | *2.85* | ✓ | ✓ | ✓ | | ✓ | |
| *6.21* | *2.85* | ✓ | ✓ | | ✓ | ✓ | |
| *6.80* | *3.04* | | ✓ | ✓ | ✓ | ✓ | ✓ |
| *7.70* | *2.96* | | ✓ | ✓ | ✓ | ✓ | |
| *8.20* | *2.90* | | ✓ | | ✓ | ✓ | ✓ |
| *9.04* | *2.95* | ✓ | ✓ | | ✓ | ✓ | ✓ |
| *9.05* | *2.85* | ✓ | ✓ | | | ✓ | ✓ |
| *9.91* | *2.98* | ✓ | ✓ | ✓ | ✓ | ✓ | |
| *10.76* | *2.88* | | ✓ | ✓ | | ✓ | ✓ |
| *38.52* | *13.42* | ✓ | ✓ | | | | |
| *38.61* | *13.29* | ✓ | ✓ | | | | ✓ |
| *39.19* | *13.38* | | ✓ | ✓ | ✓ | | ✓ |
| *39.20* | *13.37* | ✓ | ✓ | ✓ | | | |
| *39.34* | *13.43* | ✓ | ✓ | ✓ | ✓ | | ✓ |
| *39.62* | *13.26* | | ✓ | ✓ | | | |
| *39.93* | *13.34* | | ✓ | ✓ | | | ✓ |
| *40.18* | *13.39* | ✓ | ✓ | ✓ | | | ✓ |
| *40.33* | *13.40* | ✓ | ✓ | | ✓ | | ✓ |
| *40.55* | *13.31* | ✓ | ✓ | | ✓ | | |
| *40.87* | *13.30* | | ✓ | | | | |
| *41.02* | *13.53* | | ✓ | | ✓ | | ✓ |
| *41.40* | *13.34* | | ✓ | ✓ | ✓ | | |

*(continued)*

| | | λ | | | τ | | |
|---|---|---|---|---|---|---|---|
| ΔLOOIC | SE | $l_{gc}$ | $g_i$ | $g_{max}$ | $l_{gc}$ | $g_i$ | $g_{max}$ |
| 43.99 | 13.30 | ✓ | ✓ | ✓ | ✓ | | |
| 44.44 | 13.42 | | ✓ | | | | ✓ |
| 45.04 | 13.23 | | ✓ | | ✓ | | |
| 53.96 | 15.47 | | | ✓ | ✓ | ✓ | |
| 55.21 | 15.36 | | | ✓ | | ✓ | |
| 56.20 | 15.41 | ✓ | | ✓ | | ✓ | |
| 57.58 | 15.46 | | | ✓ | ✓ | ✓ | ✓ |
| 57.85 | 15.37 | ✓ | | | ✓ | ✓ | ✓ |
| 57.94 | 15.52 | ✓ | | | ✓ | ✓ | |
| 58.12 | 15.40 | ✓ | | ✓ | ✓ | ✓ | ✓ |
| 59.08 | 15.46 | ✓ | | ✓ | | ✓ | ✓ |
| 59.45 | 15.41 | | | | | ✓ | |
| 59.87 | 15.42 | ✓ | | | | ✓ | |
| 60.11 | 15.43 | | | ✓ | | ✓ | ✓ |
| 60.21 | 15.45 | ✓ | | | | ✓ | ✓ |
| 60.63 | 15.41 | | | | | ✓ | ✓ |
| 61.45 | 15.37 | | | | ✓ | ✓ | |
| 61.86 | 15.49 | | | | ✓ | ✓ | ✓ |
| 63.84 | 15.41 | ✓ | | ✓ | ✓ | ✓ | |
| 92.74 | 20.59 | | | | | | |
| 93.78 | 20.53 | | | ✓ | | | ✓ |
| 94.61 | 20.53 | ✓ | | ✓ | | | |
| 94.93 | 20.57 | | | ✓ | ✓ | | ✓ |
| 95.18 | 20.58 | ✓ | | ✓ | ✓ | | |
| 96.21 | 20.59 | | | ✓ | | | |
| 96.69 | 20.51 | ✓ | | | | | |
| 96.75 | 20.54 | ✓ | | ✓ | | | ✓ |
| 96.87 | 20.64 | | | ✓ | ✓ | | |
| 97.11 | 20.50 | | | | ✓ | | |
| 97.98 | 20.56 | ✓ | | | | | ✓ |
| 98.01 | 20.53 | ✓ | | | ✓ | | ✓ |
| 98.27 | 20.45 | ✓ | | | ✓ | | |
| 99.69 | 20.68 | ✓ | | ✓ | ✓ | | ✓ |
| 100.41 | 20.64 | | | | ✓ | | ✓ |
| 101.31 | 20.43 | | | | | | ✓ |

Table S5: Pareto $\hat{k}$, PSIS effective sample size (ESS), and Monte Carlo standard error (MCSE) diagnostics for the six plausible models identified by LOOIC (Table S4). The selected model is in bold. Good/bad/very bad refer to the conventional Pareto $\hat{k}$ thresholds of 0.7 and 1. MCSE($elpd_{loo}$) is left blank ('–') for models with at least one curve with $\hat{k} \geq 1$, for which the MCSE estimate is undefined.

| Model | ΔLOOIC | SE | $n$ curves | Good $\hat{k}$ (%) | Bad $\hat{k}$ (%) | Very bad $\hat{k}$ (%) | Min. PSIS ESS | MCSE($elpd_{loo}$) |
|---|---|---|---|---|---|---|---|---|
| **model 46** | 0.00 | 0.00 | 2125 | 99.95 | 0.05 | 0.00 | 155 | 0.99 |
| model 01 | 3.58 | 2.95 | 2125 | 99.95 | 0.05 | 0.00 | 149 | 0.99 |
| model 14 | 5.20 | 2.72 | 2125 | 100.00 | 0.00 | 0.00 | 137 | 1.00 |
| model 30 | 4.92 | 2.90 | 2125 | 99.95 | 0.05 | 0.00 | 158 | 1.00 |
| model 42 | 4.91 | 2.92 | 2125 | 100.00 | 0.00 | 0.00 | 94 | 1.00 |
| model 44 | 4.80 | 2.96 | 2125 | 100.00 | 0.00 | 0.00 | 172 | 0.99 |

Table S6: The relative contribution of phylogenetic, among population (nonphylogenetic), and among-individual varation differs among stomatal traits. The phylogenetic component is equivalent to the phylogenetic heritability. The population component is the non-phylogenetic variance among populations. The among-individual variance component is the variation among individuals within a population after accounting for treatment effect. Estimates and 95% confidence intervals (CI) are estimated as the median and quantile intervals of the posterior distribution. $l_{gc}$: guard cell length; $g_i$: initial stomatal conductance; $g_{max}$: anatomical maximum stomatal conductance; $\lambda$: lag time; $\tau$: time constant.

| component | % variance | 95% CI |
|---|---|---|
| $\log(l_{gc})$ | | |
| phylogenetic | 61.9% | [18.9%, 82.8%] |
| population (nonphylogenetic) | 7.5% | [0.6%, 36.2%] |
| among-individual | 29.9% | [15.7%, 49.2%] |
| $\log(g_i)$ | | |
| phylogenetic | 16.9% | [1.1%, 37.9%] |
| population (nonphylogenetic) | 5.6% | [0.9%, 17.2%] |
| among-individual | 77.0% | [57.7%, 88.0%] |
| $\log(g_{max})$ | | |
| phylogenetic | 59.1% | [34.4%, 75.5%] |
| population (nonphylogenetic) | 1.3% | [0.0%, 13.6%] |
| among-individual | 38.8% | [23.4%, 55.7%] |
| $\log(\lambda)$ | | |
| phylogenetic | 16.5% | [0.2%, 41.8%] |
| population (nonphylogenetic) | 4.7% | [0.2%, 15.8%] |
| among-individual | 78.3% | [56.3%, 90.3%] |
| $\log(\tau)$ | | |
| phylogenetic | 39.8% | [18.7%, 60.4%] |
| population (nonphylogenetic) | 1.1% | [0.0%, 11.2%] |
| among-individual | 58.5% | [38.7%, 74.1%] |

Table S7: Fixed effect parameter estimates and 95% confidence intervals (CIs) from the posterior distribution of the selected model. For each response variable, we estimated effects of growth light intensity (sun vs. shade), measurement light intensity (high vs. low), and leaf type (amphi vs. pseudohypo). The selected model potentially includes effects of $l_{gc}$, $g_i$, and $g_{max}$ on $\tau$ and $\lambda$.

| **Parameter** | **Estimate [95% CI]** |
|---|---|
| $\log(l_{gc})$ | |
| intercept (shade, low light, amphi) | 2.96 [2.80, 3.11] |
| effect of sun growth treatment on $\log(l_{gc})$ | 0.11 [0.11, 0.12] |
| effect of pseudohypo leaf type on $\log(l_{gc})$ | -0.06 [-0.07, -0.05] |
| $\log(g_i)$ | |
| intercept (shade, low light, amphi) | -1.71 [-1.93, -1.47] |
| effect of sun growth treatment on $\log(g_i)$ | 0.35 [0.30, 0.40] |
| effect of high measurement light intensity on $\log(g_i)$ | 0.76 [0.73, 0.78] |
| effect of pseudohypo leaf type on $\log(g_i)$ | -0.22 [-0.24, -0.19] |
| $\log(g_{max})$ | |
| intercept (shade, low light, amphi) | 0.86 [0.55, 1.17] |
| effect of sun growth treatment on $\log(g_{max})$ | 0.61 [0.59, 0.63] |
| effect of pseudohypo leaf type on $\log(g_{max})$ | -0.40 [-0.42, -0.38] |
| $\log(\lambda)$ | |
| intercept (shade, low light, amphi) | 0.39 [0.28, 0.48] |
| effect of sun growth treatment on $\log(\lambda)$ | 0.03 [0.01, 0.06] |
| effect of high measurement light intensity on $\log(\lambda)$ | -0.02 [-0.04, 0.01] |
| effect of pseudohypo leaf type on $\log(\lambda)$ | 0.01 [-0.00, 0.03] |
| effect of $\log(g_i)$ on $\log(\lambda)$ | 0.15 [0.11, 0.19] |
| $\log(\tau)$ | |
| intercept (shade, low light, amphi) | 5.58 [5.24, 5.91] |
| effect of sun growth treatment on $\log(\tau)$ | -0.11 [-0.16, -0.06] |
| effect of high measurement light intensity on $\log(\tau)$ | 0.08 [0.01, 0.15] |
| effect of pseudohypo leaf type on $\log(\tau)$ | 0.08 [0.05, 0.11] |
| effect of $\log(g_i)$ on $\log(\tau)$ | 0.32 [0.23, 0.41] |

Table S8: Raw data file in CSV format containing estimates of stomatal anatomy and kinetic variables associated with each curve. These are raw data for fitting multiresponse models. A final version will be deposited on Dryad after acceptance. This version is available for reviewers.

**Table file:**
Download from:
https://github.com/cdmuir/solanum-kinetics/raw/refs/heads/main/tables/tbl-estimates-curve.csv

Table S9: Raw data file in CSV format containing estimates of stomatal anatomy and kinetic variables associated with each population based on multiresponse model predictions. A final version will be deposited on Dryad after acceptance. This version is available for reviewers.

**Table file:**
Download from:
https://github.com/cdmuir/solanum-kinetics/raw/refs/heads/main/tables/tbl-estimates-accession.csv

Table S10: Random effect parameter estimates and 95% confidence intervals (CIs) from the posterior distribution of the selected model. For each response variable, we estimated among-individual random effects from repeated measures of the same individual, where possible, random effects of population (nonphylogenetic), and phylogenetically structured random effects of population. We report the random effect standard deviations (SD). For population-level random effects, we also estimated correlations between traits.

| **Parameter** | **Estimate [95% CI]** |
|---|---|
| *among-individual random effects* | |
| SD in $\log(\lambda)$ | 0.07 [0.07, 0.08] |
| SD in $\log(\tau)$ | 0.21 [0.19, 0.22] |
| SD in $\log(g_i)$ | 0.27 [0.25, 0.30] |
| *phylogenetic random effects* | |
| SD in $\log(\lambda)$ | 0.06 [0.01, 0.12] |
| SD in $\log(\tau)$ | 0.25 [0.16, 0.38] |
| SD in $\log(g_i)$ | 0.18 [0.05, 0.32] |
| SD in $\log(g_{max})$ | 0.27 [0.18, 0.40] |
| SD in $\log(l_{gc})$ | 0.13 [0.06, 0.21] |
| correlation between in $\log(\lambda)$ and $\log(\tau)$ | 0.42 [-0.26, 0.84] |
| correlation between in $\log(\lambda)$ and $\log(g_i)$ | -0.06 [-0.65, 0.64] |
| correlation between in $\log(\lambda)$ and $\log(g_{max})$ | -0.27 [-0.71, 0.45] |
| correlation between in $\log(\lambda)$ and $\log(l_{gc})$ | 0.09 [-0.53, 0.70] |
| correlation between in $\log(\tau)$ and $\log(g_i)$ | 0.13 [-0.49, 0.64] |
| correlation between in $\log(\tau)$ and $\log(g_{max})$ | -0.07 [-0.52, 0.37] |
| correlation between in $\log(\tau)$ and $\log(l_{gc})$ | 0.62 [0.14, 0.88] |
| correlation between in $\log(g_i)$ and $\log(g_{max})$ | 0.38 [-0.28, 0.78] |
| correlation between in $\log(l_{gc})$ and $\log(g_i)$ | 0.41 [-0.27, 0.80] |
| correlation between in $\log(l_{gc})$ and $\log(g_{max})$ | -0.00 [-0.57, 0.45] |
| *population (nonphylogenetic) random effects* | |
| SD in $\log(\lambda)$ | 0.03 [0.01, 0.06] |
| SD in $\log(\tau)$ | 0.04 [0.00, 0.13] |
| SD in $\log(g_i)$ | 0.11 [0.05, 0.19] |
| SD in $\log(g_{max})$ | 0.04 [0.00, 0.12] |
| SD in $\log(l_{gc})$ | 0.05 [0.02, 0.09] |
| correlation between in $\log(\lambda)$ and $\log(\tau)$ | 0.17 [-0.63, 0.79] |
| correlation between in $\log(\lambda)$ and $\log(g_i)$ | -0.35 [-0.83, 0.37] |
| correlation between in $\log(\lambda)$ and $\log(g_{max})$ | -0.13 [-0.76, 0.71] |
| correlation between in $\log(\lambda)$ and $\log(l_{gc})$ | -0.01 [-0.65, 0.66] |
| correlation between in $\log(\tau)$ and $\log(g_i)$ | 0.09 [-0.65, 0.74] |
| correlation between in $\log(\tau)$ and $\log(g_{max})$ | -0.06 [-0.78, 0.71] |
| correlation between in $\log(\tau)$ and $\log(l_{gc})$ | 0.24 [-0.64, 0.84] |

*(continued)*

| **Parameter** | **Estimate [95% CI]** |
|---|---|
| correlation between in log $(g_{\mathrm{i}})$ and log $(g_{\mathrm{max}})$ | -0.17 [-0.78, 0.56] |
| correlation between in log $(l_{\mathrm{gc}})$ and log $(g_{\mathrm{i}})$ | 0.44 [-0.18, 0.84] |
| correlation between in log $(l_{\mathrm{gc}})$ and log $(g_{\mathrm{max}})$ | 0.03 [-0.72, 0.71] |

### Supporting figures

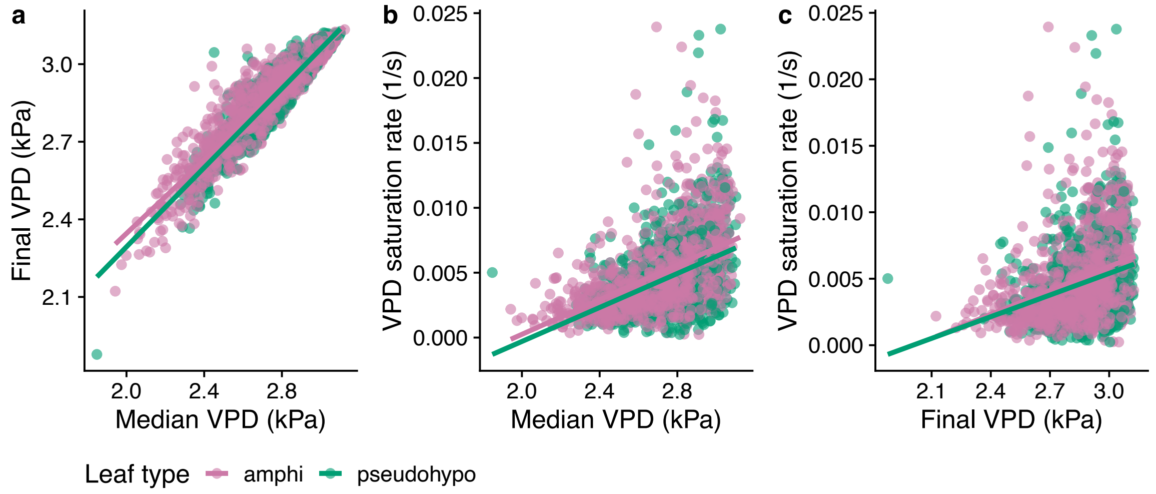


Figure S1: **Pairwise relationships among curve-level VPD covariates. a.** Median vs. final VPD during the fitted interval; **b.** median VPD vs. VPD saturation rate $k$; **c.** final VPD vs. VPD saturation rate $k$. Each point is one humidity response curve, colored by leaf type; lines are ordinary least-squares fits within leaf type. VPD saturation rate is the rate constant of a saturating exponential fit to the VPD trajectory during the fitted interval (see Notes S1).

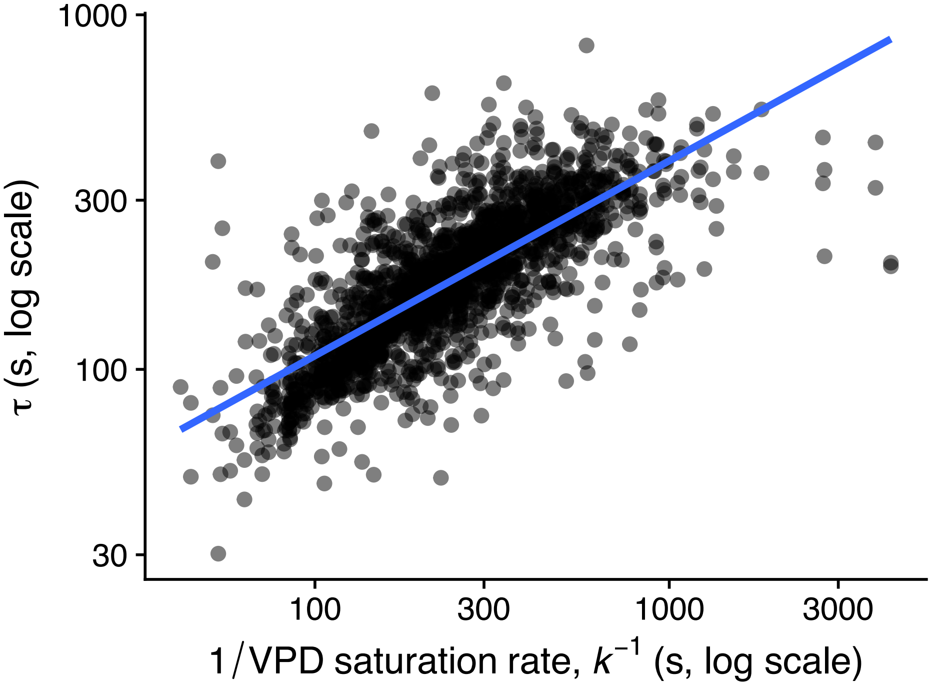


Figure S2: **The stomatal closure time constant $\tau$ (log scale) is strongly correlated with the inverse of the VPD saturation rate $k^{-1}$ ($r = 0.77$ on log-log scale)**. This is consistent with $k$ being driven largely by the same $g_{sw}$-decline signal that determines $\tau$, rather than an independent property of the VPD stimulus (see Notes S1). Each point is one humidity response curve from the untreated (amphi) leaves. The blue line us ordinary least-squares regression fit.

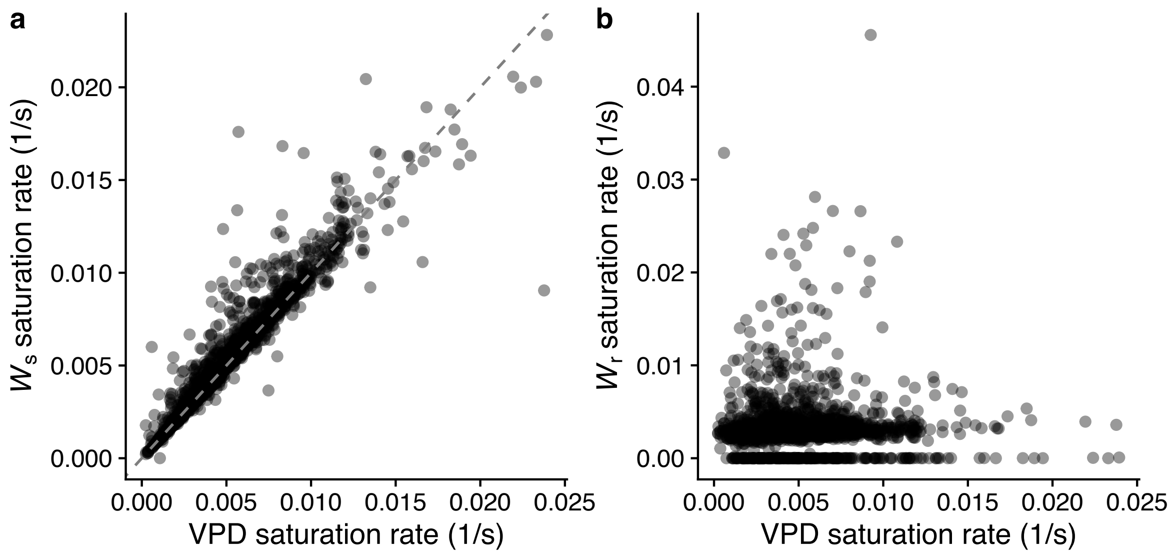


Figure S3: **The VPD saturation rate is driven by the sample airstream, not the reference airstream.** **a.** VPD saturation rate vs. the saturation rate of $W_s$ (sample cell water vapor concentration); the dashed line is the 1:1 line. **b.** VPD saturation rate vs. the saturation rate of $W_r$ (reference cell water vapor concentration, upstream of the leaf). Each point is one humidity response curve; rate constants come from the same saturating-exponential model used for VPD.

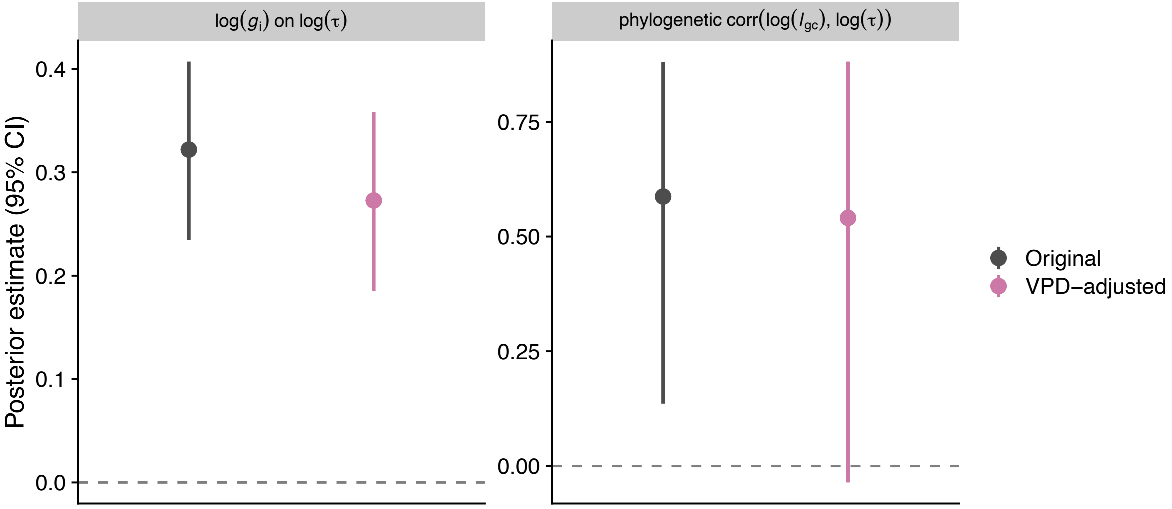


Figure S4: **The $g_i$ effect on $\tau$ and the phylogenetic correlation between $l_{gc}$ and $\tau$ are robust to accounting for final VPD.** Posterior means and 95% credible intervals for the fixed effect of $f_{gmax}$ on $\log \tau$ (left) and the phylogenetic correlation between $\log l_{gc}$ and $\log \tau$ (right), comparing the original model (grey) to a model refit with final VPD added as a covariate of $\tau$ and $\lambda$ (pink; see Notes S1).

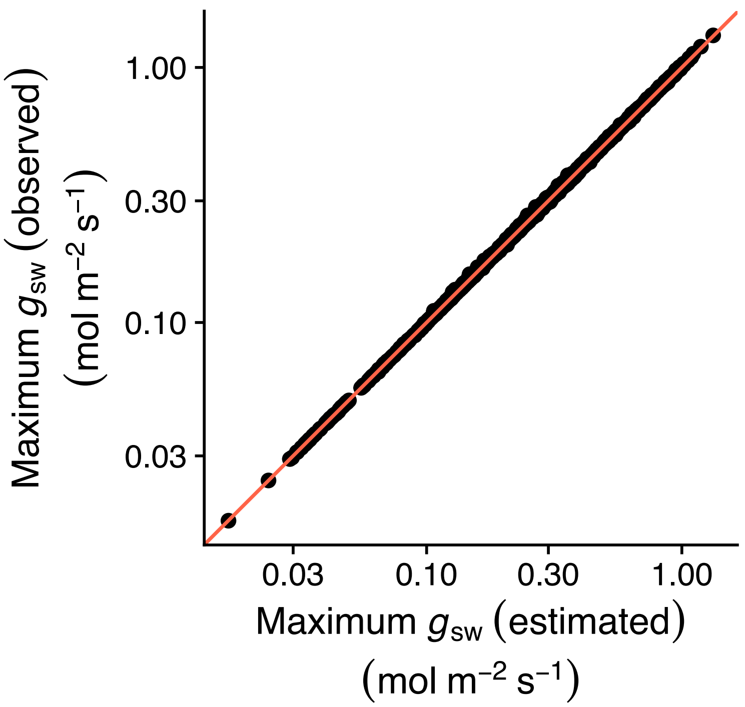


Figure S5: **The estimated ($x$-axis) and observed ($y$-axis) maximum stomatal conductances ($g_{sw}$) are nearly identical.** Each point is the empirical maximum $g_{sw}$ plotted against the estimated $g_{sw}$ for every humidity response curve. The orange line is the 1:1 line for reference. Both axes are on a log-scale.

**Figure file:**
Download from:
https://github.com/cdmuir/solanum-kinetics/raw/refs/heads/main/figures/rh-curves.pdf

Figure S6: **Humidity-response curves and fitted lines.** The title provides the Tomato Genetics Resource Center accession number, replicate letter, and species names. The subtitle indicates the growth light intensity (sun or shade), measurement light intensity (low or high), and leaf type (amphi or pseudohypo). Points are raw data and lines are fitted curves. The Bayesian correlation coefficient, Bayes $R^2$, is shown to the right of each curve.

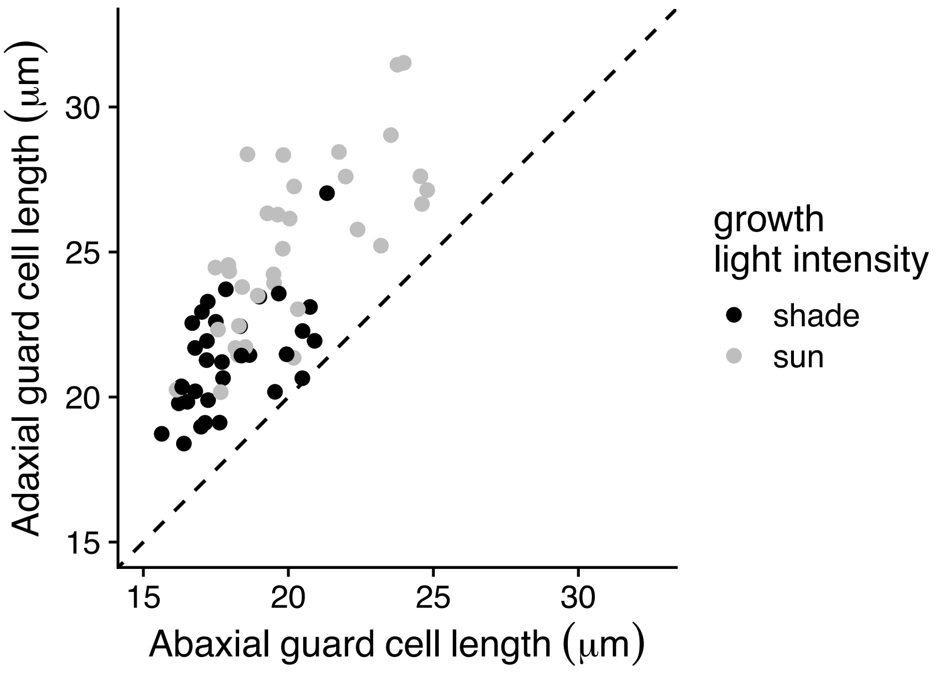


Figure S7: **Adaxial (upper) stomata are consistently larger than abaxial (lower) stomata in both sun-grown (grey points) and shade-grown (black points) plants.** Each point was calculated by taking the median guard cell length within each biological replicate and then calculating the mean among replicates within each population and growth light intensity treatment. The dashed line is the 1:1 line for reference. All points above this line indicate larger values on the adaxial surface.

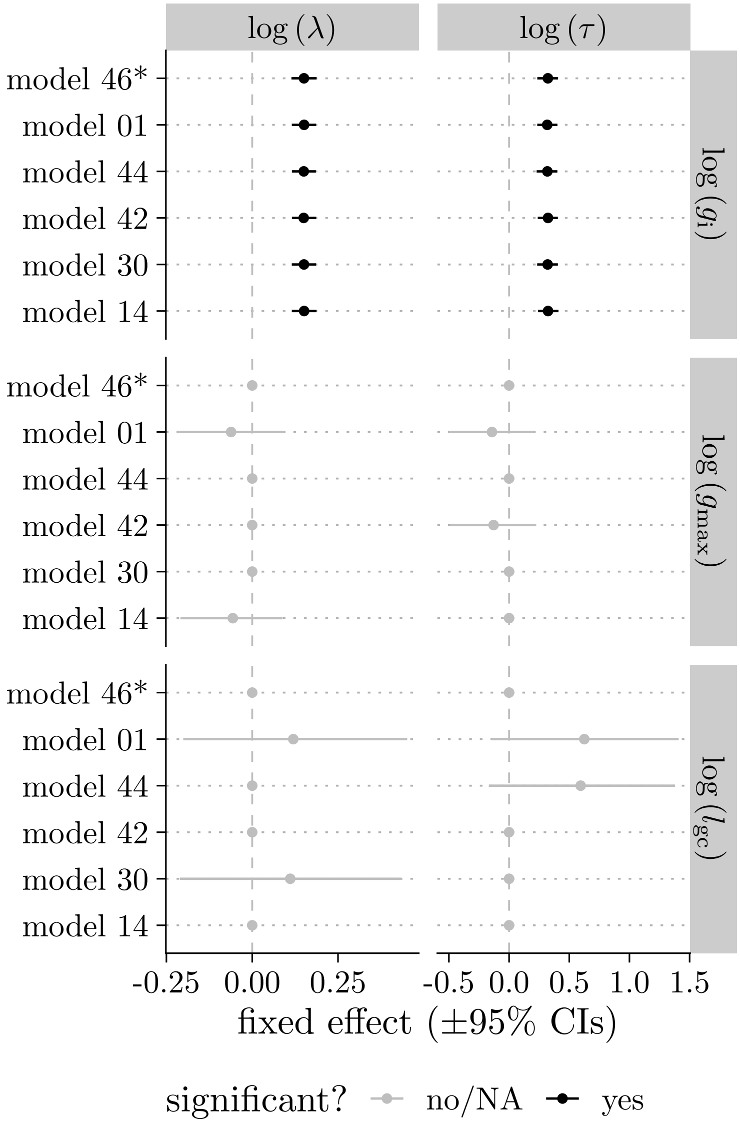


Figure S8: **Individual-level (fixed-effect) estimates of initial stomatal conductance ($g_i$, top facets), anatomical maximum stomatal conductance ($g_{max}$, middle facets), and guard cell length ($l_{gc}$, bottom facets) on the stomatal closure lag time ($\lambda$, left facets) and time constant ($\tau$, right facets) are broadly consistent across all plausible models.** Points and horizontal bars are posterior medians and 95% credible intervals from each of the six plausible models (rows; Table S4), ordered by ΔLOOIC; the selected model is marked with an asterisk (*). Black indicates estimates whose 95% credible interval excludes zero; grey indicates intervals that overlap zero.

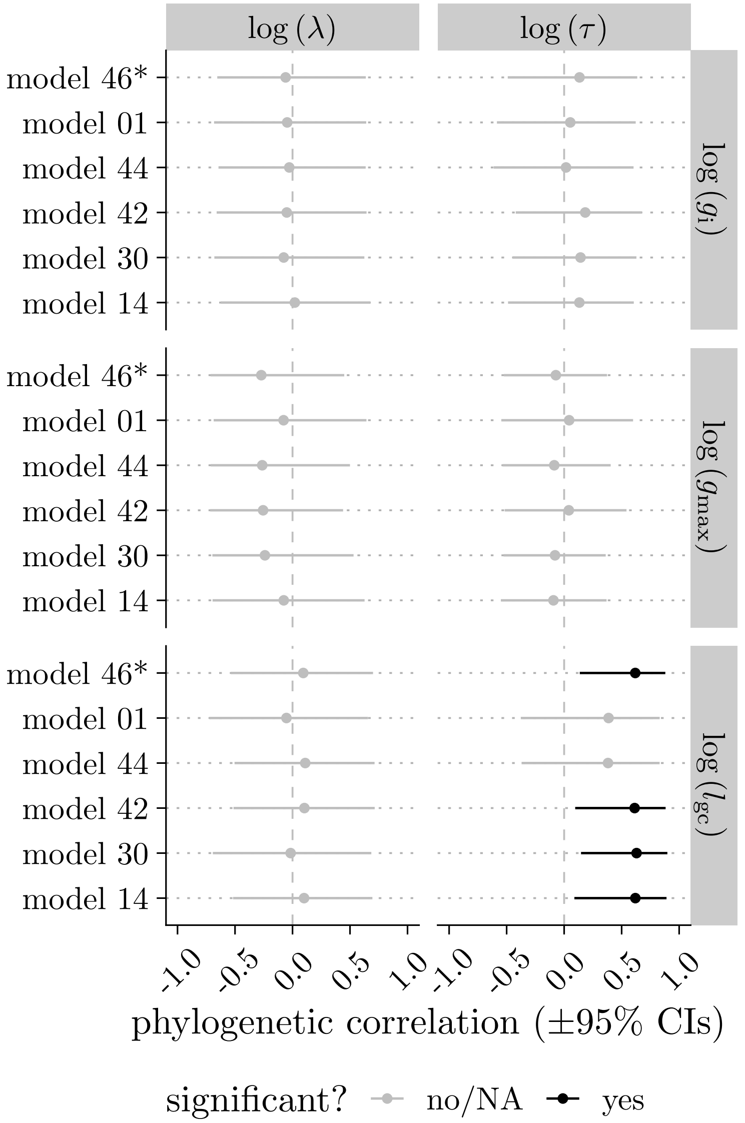


Figure S9: **Phylogenetic correlations between initial stomatal conductance ($g_i$, top facets), anatomical maximum stomatal conductance ($g_{max}$, middle facets), and guard cell length ($l_{gc}$, bottom facets) on the stomatal closure lag time ($\lambda$, left facets) and time constant ($\tau$, right facets) are broadly consistent across all plausible models.** Points and horizontal bars are posterior medians and 95% credible intervals of the phylogenetic correlation from each of the six plausible models (rows; Table S4), ordered by ΔLOOIC; the selected model is marked with an asterisk (*). Black indicates estimates whose 95% credible interval excludes zero; grey indicates intervals that overlap zero.

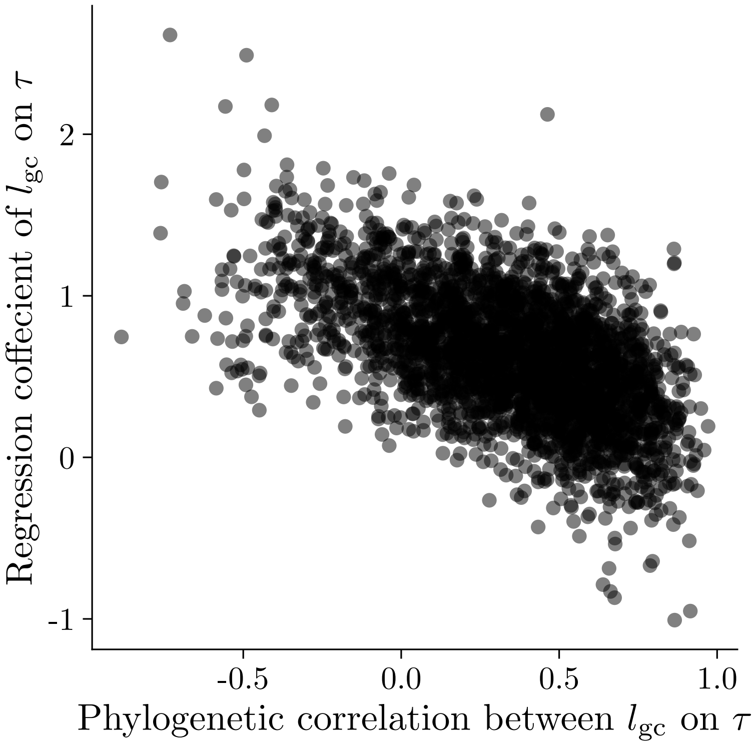


Figure S10: **Posterior estimates of phylogenetic correlations ($x$-axis) and fixed effects ($y$-axis) of guard cell length ($l_{\mathrm{gc}}$) on the time constant ($\tau$) are collinear.** Each point is the estimate from one draw of the posterior distribution for both parameters from the model with the lowest LOOIC. The negative relationship indicates that there is tradeoff in fitting the relationship between variables as a higher level phylogenetic effect versus a lower level, individual effect. This results in more uncertainty and larger confidence intervals in each individual parameter.

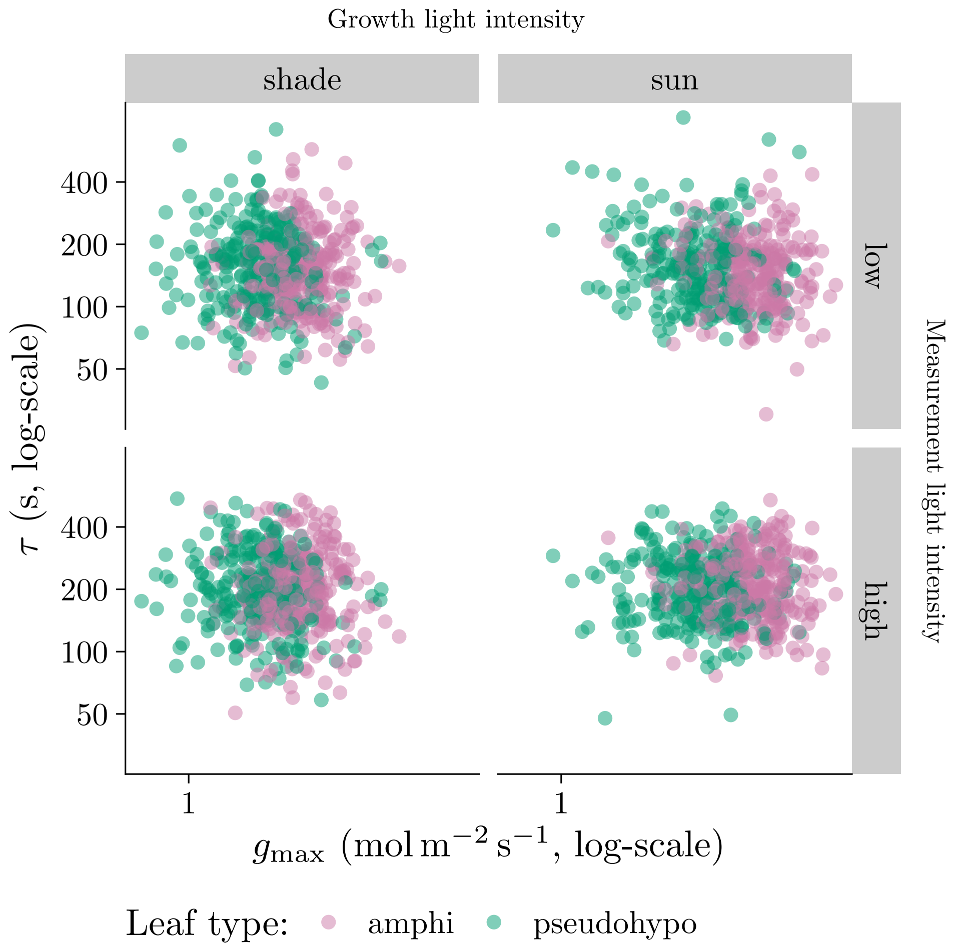


Figure S11: **Individual-level variation in anatomical maximum stomatal conductance ($g_{\text{max}}$) does not influence the stomatal closure time constant $\tau$.** Individual-level variation in $g_{\text{max}}$ ($x$-axis) did not significantly covary with $\tau$ ($y$-axis). The overall pattern was consistent across growth light intensity treatments (left and right facets), measurement light intensity treatments (top and bottom facets), and leaf types (point colors). The growth and measurement light intensity treatments are described in the Materials and Methods section.

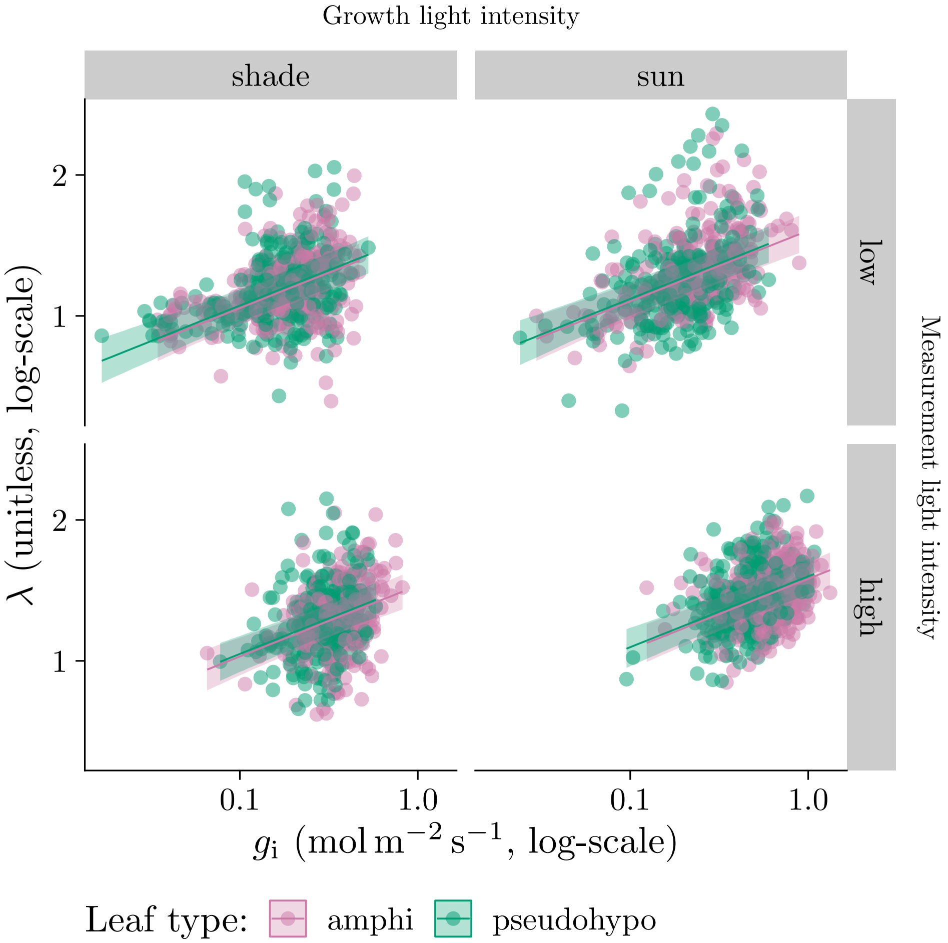


Figure S12: **Individual-level variation in initial stomatal conductance ($g_i$) influences stomatal closure kinetics.** As $g_i$ ($x$-axis) increases, the stomatal closure lag time ($\lambda$, $y$-axis) increases. The overall pattern was consistent across growth light intensity treatments (left and right facets), measurement light intensity treatments (top and bottom facets), and leaf types (point colors). Lines are estimated using linear regression along with 95% confidence ribbons. The growth and measurement light intensity treatments are described in the Materials and Methods section.

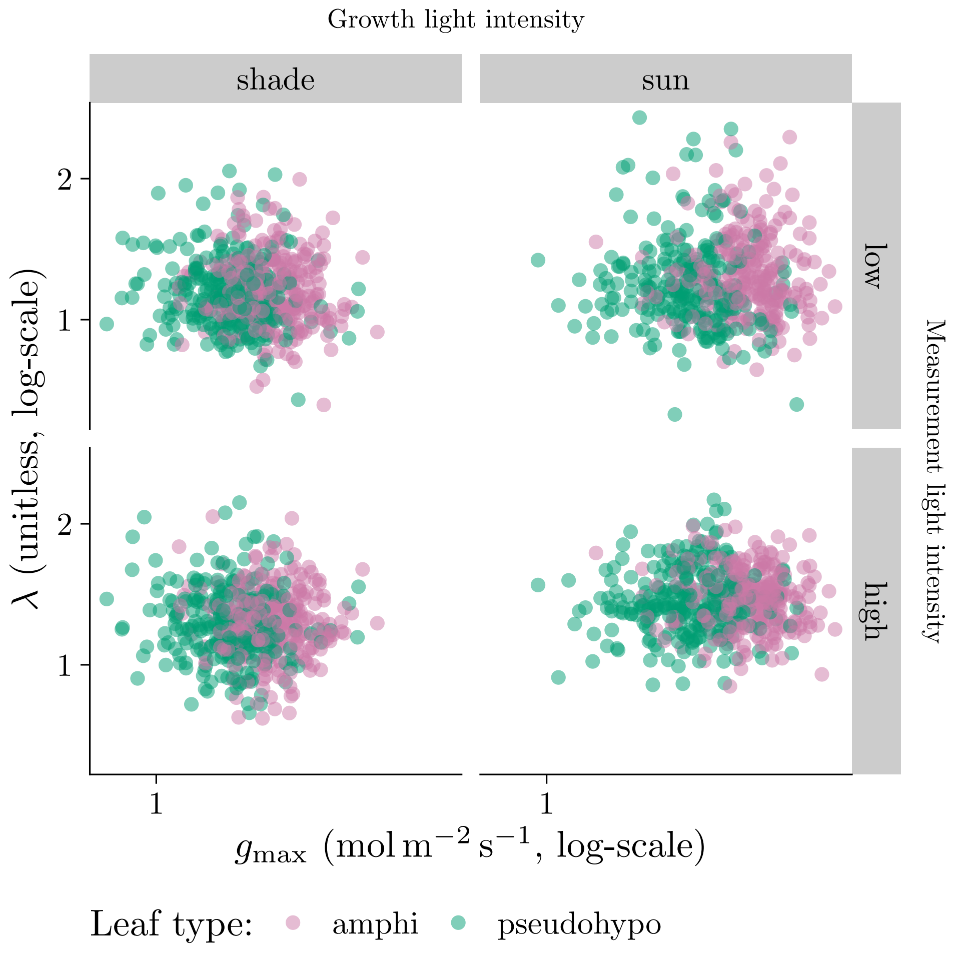


Figure S13: **Individual-level variation in anatomical maximum stomatal conductance ($g_{max}$) does not influence the stomatal closure lag time $\lambda$.** Individual-level variation in $g_{max}$ ($x$-axis) did not significantly covary with $\lambda$ ($y$-axis). The overall pattern was consistent across growth light intensity treatments (left and right facets), measurement light intensity treatments (top and bottom facets), and leaf types (point colors). The growth and measurement light intensity treatments are described in the Materials and Methods section.

### Supporting notes

#### Notes S1: Effects of $g_{\mathrm{i}}$ and $l_{\mathrm{gc}}$ are robust to variation in VPD stimulus

Table S2 shows that the mean VPD realized during measurements differed among treatment combinations (2.72 kPa–3 kPa), because leaf transpiration itself humidifies the chamber and $g_{\mathrm{sw}}$ varied systematically with measurement light, growth light, and leaf type. Because VPD is the physiological driver of stomatal closure, we checked whether this variation in the realized stimulus could confound the effects of $g_{\mathrm{i}}$ and $l_{\mathrm{gc}}$ on the closure time constant, $\tau$.

For every curve we extracted the leaf-to-air VPD trajectory from the logged humidity data and derived two curve-level covariates: final VPD and VPD saturation rate ($k$). Final VPD is the VPD at the end of the fitted interval; $k$ is the rate constant of a saturating exponential, $\text{VPD}(t) = B + (R_0 - B)\exp(-kt)$, fit separately to each curve (median $R^2 = 0.99$, $n = 2,156$ curves; $k$ excluded for 15 curves with a poor saturating fit or an extreme outlier rate estimate). We also considered median VPD. Median and final VPD were highly correlated ($r = 0.93$; Figure S1a), so we retained only final VPD.

We did not include the VPD saturation rate as a covariate of $\tau$, because $k$ and $\tau$ are themselves strongly correlated ($r = 0.77$; Figure S2) for a mechanistic rather than confounding reason: the VPD trajectory during the analyzed interval is driven mainly by declining leaf transpiration as stomata close, so $k$ largely tracks the same $g_{\mathrm{sw}}$-decline signal that determines $\tau$. We tested this by fitting the same saturating-exponential model to the sample-cell water vapor concentration ($W_{\mathrm{s}}$) and the reference-cell water vapor concentration ($W_{\mathrm{r}}$, upstream of the leaf and largely reflecting the controlled incoming airstream). The VPD saturation rate closely tracked the $W_{\mathrm{s}}$ saturation rate ($r = 0.95$) but not the $W_{\mathrm{r}}$ saturation rate ($r = 0.02$) (Figure S3). Including $k$ as a covariate of $\tau$ would therefore be close to circular.

To directly test whether the realized VPD stimulus confounds the $g_{\mathrm{i}}$ and $l_{\mathrm{gc}}$ effects on $\tau$, we refit the selected model with final VPD added as a covariate of $\tau$ and $\lambda$ (Figure S4). The individual-level effect of $g_{\mathrm{i}}$ on $\log(\tau)$ was slightly weaker (original: 0.32 [0.23, 0.41]; VPD-adjusted: 0.27 [0.18, 0.36]), as was the phylogenetic correlation between $l_{\mathrm{gc}}$ and $\tau$ (original: 0.59 [0.14, 0.88]; VPD-adjusted: 0.54 [-0.04, 0.88]). This indicates that variation in VPD slightly amplified the apparent effects of $g_{\mathrm{i}}$ and $l_{\mathrm{gc}}$ on stomatal closure kinetics, but does not qualitatively change the main conclusions. Final VPD itself had a negative fixed effect on $\log(\tau)$ (-0.41 [-0.53, -0.27]), consistent with higher realized VPD accelerating stomatal closure.

We lack data on the rate of VPD change during the wrong-way response and the earliest part of the right-way response, because humidity logging did not begin until $g_{\mathrm{sw}}$ had already declined back to its pre-step value (Figure 1a); this limits our ability to evaluate whether the *initial* rate of VPD change, as opposed to the final realized VPD, influenced closure kinetics. We recommend that future studies of stomatal kinetics log chamber humidity continuously from the onset of the humidity step to more fully characterize the realized VPD stimulus and adjust the incoming airstream to keep VPD constant as stomatal close.

**Notes S2: The $g_i$-$\tau$ association is not an artifact of the curve-fitting procedure**

The initial conductance $g_i$ used to calculate $f_{gmax}$ is estimated from the same nonlinear fit (Equation 1) that also estimates $\tau$ and $\lambda$ for each curve. This raises the possibility that parameter covariance within the curve fit itself, rather than a true biological relationship, could generate an association between $g_i$ and $\tau$ (and hence $f_{gmax}$ and $\tau$, since the denominator $g_{max}$ is estimated independently from stomatal anatomy and cannot itself introduce a fitting artifact). To test this, we ran a null simulation in which any true association between $g_i/g_f$ and the kinetic parameters was removed by construction. For every real curve $i$, we simulated a synthetic $g_{sw}$ trajectory using curve $i$'s own measurement times, residual noise magnitude, and fitted $\tau$ and $\lambda$, but with $g_i$ and $g_f$ taken from a different, randomly chosen curve $j$. Because $i$ and $j$ were chosen independently at random, the synthetic curves retain realistic variability in both $\tau/\lambda$ and $g_i/g_f$, but there is no true association between $g_i$ and the kinetic parameters. We re-fit the same Weibull functional form to each synthetic curve using nonlinear least squares (`nls()`), which closely approximated the full Bayesian pipeline used for the real data when compared directly on the real curves (cor($\tau$) = 0.99, cor($\lambda$) = 1.00), and computed the correlation between the *estimated* $g_i$ and $\log(\tau)$. We repeated this 1,000 times to build a null distribution of this correlation.

The null-simulation correlations ranged from -0.08 to 0.08 (median $-3 \times 10^{-3}$) across the 1,000 replicates, indistinguishable from zero (Figure S14). In contrast, the real, observed correlation between $g_i$ and $\log(\tau)$ is 0.43 [0.39, 0.46], far outside the null range (empirical $p = 0.000$). This indicates that the curve-fitting procedure alone cannot explain the observed $g_i$-$\tau$ association, supporting our interpretation that it reflects a real relationship in the data rather than a statistical artifact of estimating $g_i$, $\tau$, and $\lambda$ from the same nonlinear fit.

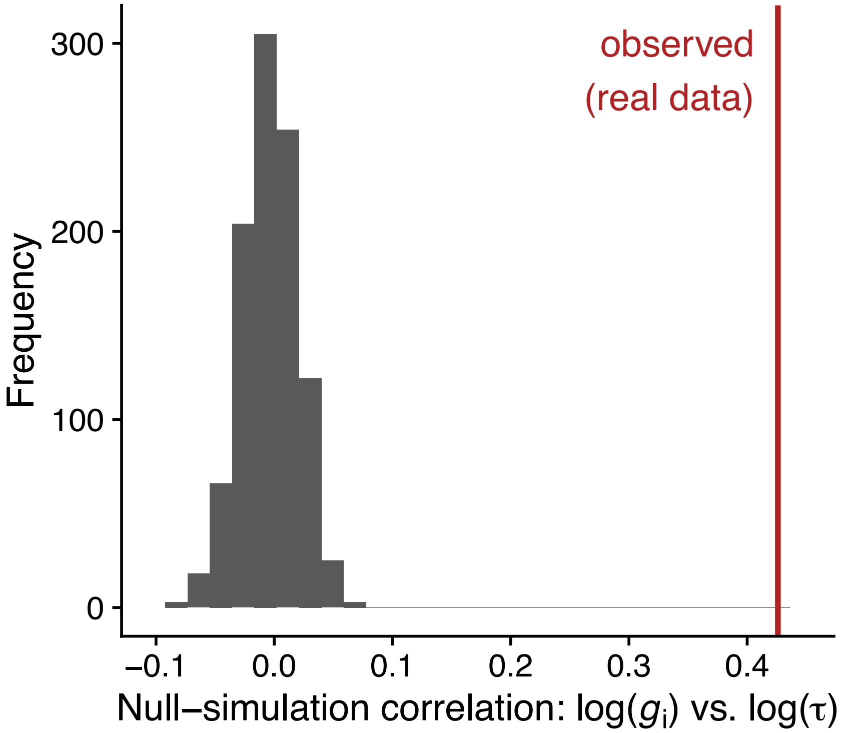


Figure S14: **The curve-fitting procedure alone does not induce a $g_{\mathrm{i}}$-$\tau$ association.** Null distribution of the correlation between estimated $g_{\mathrm{i}}$ and estimated $\tau$ across 1,000 simulated datasets in which the true $g_{\mathrm{i}}$ and $g_{\mathrm{f}}$ are fixed at their real median values for every curve (so $g_{\mathrm{i}}$ has no true relationship with $\tau$ by construction), compared to the observed correlation in the real data (red line).

**Notes S3: Sensitivity analysis of the $g_i$-$\tau$ path analysis to unmeasured confounding**

The path analysis linking treatment, $g_i$, and $\tau$ assumes the causal structure in Figure S15: each treatment has a direct effect on $\tau$ and an indirect effect acting through $g_i$. This structure is only identified as causal under sequential ignorability, i.e., that no unmeasured confounder $U$ (e.g., hydraulic status, measurement order, realized VPD trajectory) affects both $g_i$ and $\tau$. If such a $U$ exists (dashed arrows), the correlation it induces between the $g_i$ and $\tau$ equations' residuals is exactly the sensitivity parameter $\rho$ examined below.

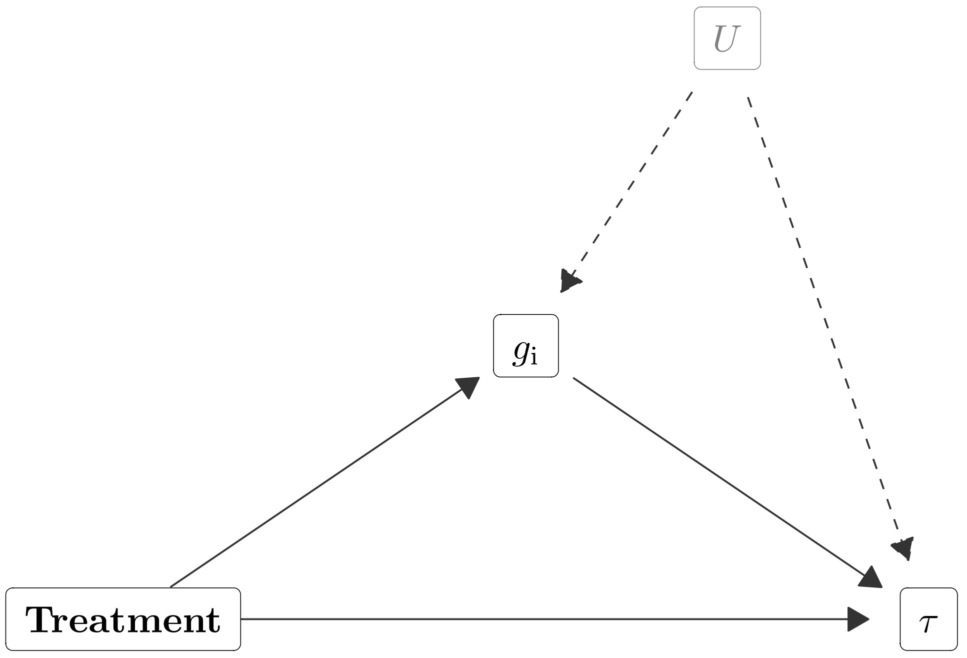


Figure S15: **Assumed causal structure of the $g_i$-$\tau$ path analysis.** Treatment has a direct effect on $\tau$ (bottom arrow) and an indirect effect acting through $g_i$ (upper path). An unmeasured confounder $U$ (dashed arrows) affecting both $g_i$ and $\tau$ – e.g., hydraulic status, measurement order, or realized VPD trajectory – would violate the sequential ignorability assumption required to interpret the indirect path causally.

The path analysis linking $g_i$ to $\tau$ (see Materials and Methods, "Path analysis to test for plastic effects") assumes sequential ignorability: that there is no unmeasured confounder of the $g_i$-$\tau$ relationship after conditioning on treatment and the random effects already in the model. Because $g_i$ is not randomized, is estimated from the same curve as $\tau$, and could share unmeasured confounders (e.g., hydraulic status, measurement order, realized VPD trajectory) with $\tau$, we assessed this assumption directly with a sensitivity analysis following the general framework of Imai *et al.* (2010), parameterized by $\rho = \mathrm{Cor}(e_{g_i}, e_\tau)$, the residual correlation between the $g_\mathrm{i}$ and $\tau$ equations. Under a violation of sequential ignorability with residual correlation $\rho$, the bias-adjusted mediator-on-outcome coefficient is $\beta_2(\rho) = \hat{\beta}_2 - \rho \cdot \sigma_\tau / \sigma_{g_i}$, so the indirect effect of a treatment acting through $g_\mathrm{i}$ is $\gamma_1 \cdot \beta_2(\rho)$ (where $\gamma_1$ is the effect of treatment on $g_\mathrm{i}$), and the "breakdown point" $\rho^*$ at which this indirect effect is exactly zero is $\rho^* = \hat{\beta}_2 \cdot \sigma_{g_i} / \sigma_\tau$.

Our selected model already estimates a free residual correlation between the $g_\mathrm{i}$ and $\tau$ equations, because we fit all five responses ($l_\mathrm{gc}$, $g_\mathrm{i}$, $g_\mathrm{max}$, $\tau$, $\lambda$) jointly. Because the $g_\mathrm{i}$ equation's predictors are a strict subset of $\tau$'s, jointly estimating this correlation does not change the point estimate of $\hat{\beta}_2$ relative to a model that assumes $\rho = 0$ (a standard result for seemingly unrelated regression models), so we can treat $\hat{\beta}_2$ as the "naive" coefficient in the sensitivity framework above, and the model's own estimated residual correlation as a data-driven estimate of $\rho$.

We found $\rho_\mathrm{estimated}$ = 0.06 [-0.07, 0.18] and $\rho^*$ = 0.40 [0.29, 0.51] (Figure S16). These posterior distributions do not substantially overlap and the posterior probability that the estimated residual correlation is smaller in magnitude than the breakdown point is nearly 100%. In other words, the plastic mediation effects of $g_\mathrm{i}$ on $\tau$ are probably not explained by unmeasured confounders.

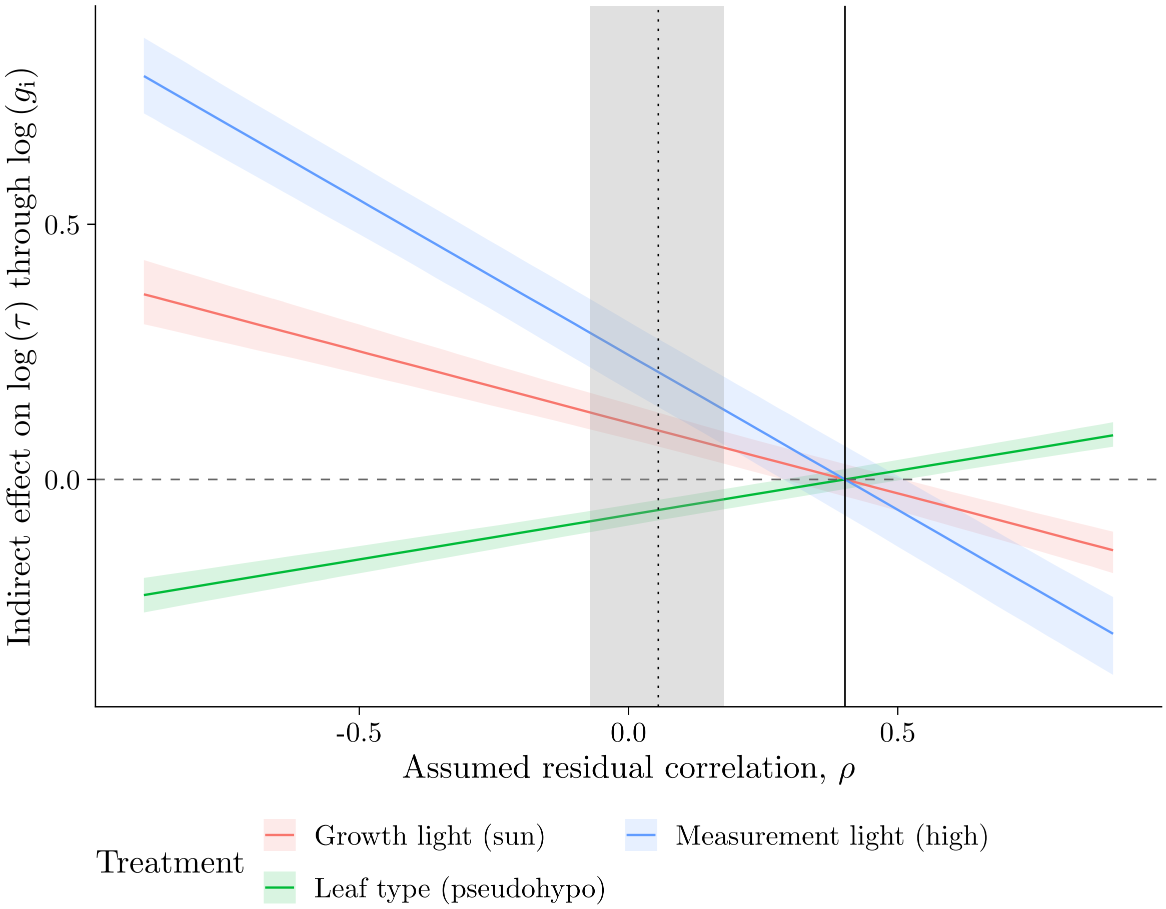


Figure S16: **Sensitivity of the indirect effect of each treatment on** $\log \tau$**, acting through** $g_\mathrm{i}$**, to an assumed residual correlation** $\rho$ **between the** $g_\mathrm{i}$ **and** $\tau$ **equations.** The vertical dotted line is the model's own estimated residual correlation ($\rho_\mathrm{estimated}$, shaded band = 95% credible interval); the vertical solid line is the breakdown point $\rho^*$ at which the indirect effect is zero.
